# Proteome-wide Profiling of RNA-Binding Protein Responses to flg22 Reveals Novel Components of Plant Immunity

**DOI:** 10.1101/2020.09.16.299701

**Authors:** Marcel Bach-Pages, Honglin Chen, Nattapong Sanguankiattichai, Riccardo Soldan, Farnusch Kaschani, Markus Kaiser, Shabaz Mohammed, Renier A.L. van der Hoorn, Alfredo Castello, Gail M. Preston

## Abstract

RNA-binding proteins (RBPs) play critical roles in post-transcriptional gene regulation and are known to contribute to plant immunity. To understand the responses of cellular RBPs to an immune elicitor, we applied RNA interactome capture to Arabidopsis leaves treated with flg22. Strikingly, flg22 induced a pervasive remodelling of the cellular RBPome affecting 186 proteins. Flg22-responsive RBPs included classical RBPs involved in RNA metabolism as well as non-canonical RBPs. RBP responders detected after 2h of treatment are enriched in putative sites for post-translational modifications, which may play a regulatory role. By contrast, changes in RBP abundance becomes increasingly important for the RBPome responses to flg22 after 12h. Plant resistance to *Pseudomonas syringae* is strongly altered in mutant lines lacking individual flg22-responsive RBPs, supporting the importance of RBP dynamics in plant immunity. This study provides a comprehensive and systematic census of flg22 responsive plant RBPs, discovering novel components of plant immunity.

## INTRODUCTION

Plants have evolved a sophisticated immune system able to effectively restrict and counteract pathogens. Plants can perceive pathogens through recognition of pathogen-associated molecular patterns (PAMPs) or damage-associated molecular patterns (DAMPs) using pattern recognition receptors (PRR)^1^. Flg22 is a 22-amino acid peptide derived from the bacterial flagella that elicits PAMP-triggered immunity (PTI) through its recognition by the plant flagellin-sensing 2 (FLS2) receptor^1^. Pathogens often evolve effector proteins able to interfere with host processes, including PTI. However, some plants can detect these effectors using nucleotide-binding site leucine-rich repeat (NBS-LRR) receptors and elicit effector-triggered immunity (ETI)^2^. Plant defence responses typically involve production of reactive oxygen species (ROS), callose deposition, inhibition of photosynthesis, stomatal closure and production of antimicrobial compounds, amongst others^3^. However, for many of these immune responses to occur, an extensive reprogramming of defence-related genes is required^4^. RNA-binding proteins (RBPs) are critical players in the post-transcriptional control of gene expression that regulate RNA processing, localization translation and stability^5^. However, it remains unknown to what extent RBPs are orchestrators of the re-programming of the transcriptome during plant immunity. Previous studies have shown the importance of several individual RBPs in defences against bacteria, fungi and other pathogens^6^. Other RBPs have been shown to be targeted by pathogen effectors, suggesting that they play important roles in immunity^6^. Although the mechanism of action of some of these RBPs is known, the scope of immunity related RBPs and their mode of action remain largely unexplored.

A proteome-wide approach known as ‘RNA interactome capture’ (RIC) was recently developed to identify cellular RBPs and study their dynamics^7,8^. RIC employs irradiation with ultraviolet (UV) light to promote crosslinks between protein and RNA interacting at ‘zero distance’, followed by capture of polyadenylated RNAs with oligo(dT) magnetic beads under denaturing conditions. RIC has dramatically increased the census of RBPs, identifying hundreds of proteins that were previously unknown to bind RNA^9^. Since then, RIC has rapidly expanded to a wide range of organisms including yeast, worm, human cell lines and plants, opening new horizons for the study of RBPs in tissues^9^. RIC has recently been used to study the dynamics of RNA-binding proteomes (RBPomes) under different experimental conditions^10,11^, offering the opportunity to investigate which RBPs respond to physiological and environmental cues as well as pathological insults. These studies have shown that the RBPome is dynamically regulated to govern the changes that occur during adaptation to physiological and environmental cues. For example, more than two hundred human RBPs differentially associate with RNA upon viral infection^10^. Moreover, the *Drosophila* RBPome changes dynamically throughout the maternal-to-zygotic transition to orchestrate post-transcriptional changes whereby the maternal mRNAs are replaced by zygotic mRNAs^11^.

Several groups have recently applied RIC to plant samples, discovering hundreds of novel plant RBPs^12–15^. Building on these advances, a recent study applied RIC to profile RBP dynamics in plant suspension cultures in response to drought stress^16^. Despite being an important first step towards the study of RBP dynamics in plants, the use of cell cultures and polyethylene glycol as a drought-mimic agent raises questions about the physiological relevance of this system. Hence, it becomes critical to adapt RIC to plant tissues to study RBP dynamics in a physiological context. However, plant tissues present several challenges for RIC studies, including the presence of cell wall and abundant pigments that absorb UV light.

Here we applied our recent plant-optimised RIC protocol (ptRIC)^12^ to leaves treated with the immunogenic peptide flg22. Strikingly, our study uncovers 186 flg22-responsive RBPs that display differential association with RNA upon immunity activation. After 2h of flg22 treatment, the majority of these changes in RNA binding cannot be explained by alterations in RBP abundance. However, the significant enrichment of post-translational modification sites in early flg22-responsive RBPs highlights the activation of signalling pathways as a potential regulatory layer of RBP activity in immunity. Later upon flg22 elicitation (12h), protein abundance becomes a driver shaping the RBPome. Through phenotypic analysis of Arabidopsis mutant lines for a subset of identified RBPs, we discovered that flg22-responsive RBPs are important for plant immunity as plants become more resistant or susceptible to infection. Taken together, we present here a comprehensive analysis of RBP dynamics in response to flg22 immune elicitation that provides new insights into RNA biology in plant immunity.

## RESULTS

### The Arabidopsis leaf RBPome and its responses to flg22

Multiple RBPs have been described to have crucial roles during plant immune responses. Hence, we hypothesised that the Arabidopsis RBPome is modulated during immune responses, and that these changes in RNA binding activity could be profiled using RIC^17^. To capture the dynamics of the plant RBPome during immune responses, Arabidopsis leaves were infiltrated with either H_2_O (mock) or the immune elicitor flg22 and leaf tissue was harvested at 2 h and 12 h post treatment (hpt). Protein-RNA interactions were ‘frozen’ by UV irradiation and RBPs covalently linked to RNA were isolated following a recently developed, plant-optimised version of RIC, referred to here as ptRIC^12^ (**Fig. 1a**,**c**). RBPs and their dynamics were then revealed by label-free quantitative proteomics.

**Fig. 1.**
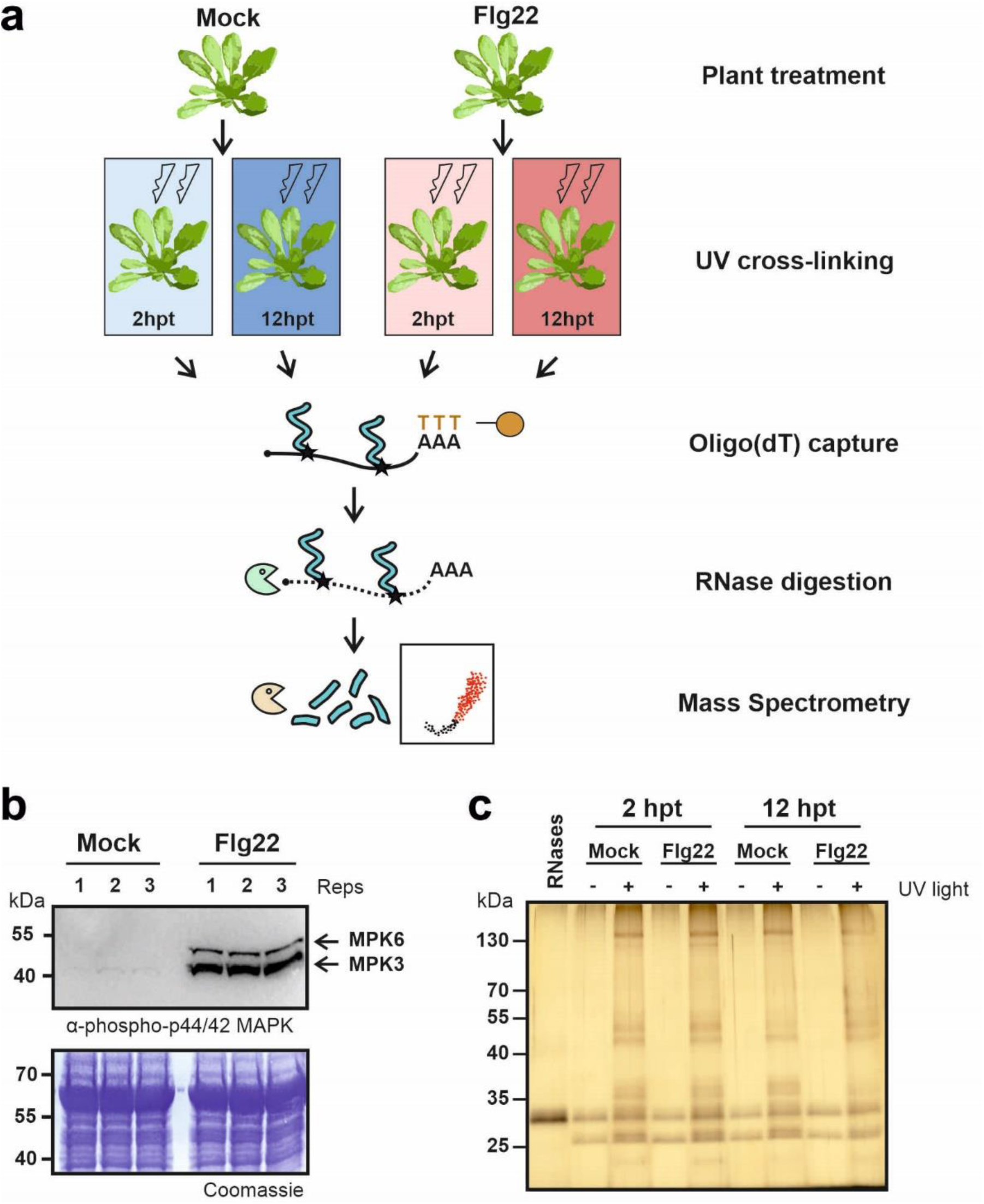
a Overview of plant RNA Interactome Capture (ptRIC) and experimental design. The leaves of mature Arabidopsis plants (5-6 weeks old) were infiltrated with either H_2_O (mock) or the immune elicitor flg22 (1 μM). The infiltrated leaves were harvested at 2 and 12 hpt and irradiated with UV light at 254 nm to promote crosslinking between RNAs and proteins that are in intimate contact. Next, cells were lysed and mRNAs captured using oligo(dT) magnetic beads. After stringent washes, the RNA-protein complexes were recovered and the RBPs released by RNase digestion. Proteins were quantitatively analysed by mass spectrometry after trypsin digestion. Four biological replicates per time point and treatment were analysed. **b Western blotting analyses of phosphorylated MAPK**. Leaves of Arabidopsis plants were infiltrated with either H_2_O or the immune elicitor flg22 (1 μM). Leaves were harvested 30 minutes after infiltrating, the proteins extracted and separated by SDS-PAGE. Proteins were transferred onto a nitrocellulose membrane and analysed by western blot using an antibody recognising phosphorylated MAPK (anti-phospho-p44/42 MAPK antibody; CST #9102). For each treatment, three biological replicates were performed (Reps 1-3). **c Silver staining analyses of the eluates (RNA-bound RBPs) isolated by ptRIC**. The RBPs isolated using ptRIC described in panel ‘**a**’ were analysed by SDS-PAGE followed by silver staining. The RNases used to release the RBPs from RNAs are loaded as a control.

To confirm that the flg22 treatment elicited plant immunity, we analysed activation of immune signalling by phosphorylation of mitogen-activated protein kinases (MAPKs)^18^. Western blotting (WB) analyses showed that MPK6 and MPK3 were phosphorylated after 30 min of flg22 treatment, which indicates that plant immunity was rapidly and efficiently elicited by flg22 (**Fig. 1b**).

Before studying the dynamics of the RBPome upon flg22 treatment, the individual leaf RBPomes for each of the 4 different conditions (mock 2 hpt, flg22 2 hpt, mock 12 hpt and flg22 12 hpt) were determined by analysing the enrichment of the proteins identified in crosslinked (CL; +UV) over not crosslinked (NoCL; -UV) samples as previously described^12^ (**Fig. S1a-d**). A total of 890 proteins were identified across all conditions in a UV-dependent manner (i.e. RBPs), constituting the ‘core’ Arabidopsis leaf RBPome (**Fig. S1e; Table S1**). However, 245 additional RBPs were identified in 3 or fewer experimental conditions, suggesting that they are state-specific RBPs (**Fig. S1e; Table S1**). The resulting dataset is compositionally similar to previously established RBPomes as 41.6% of the identified proteins are annotated by the gene ontology (GO) term ‘RNA binding’, and an additional 15.7% have annotations linked to RNA biology (**Fig. S1f**). In addition, from the 1,135 RBPs identified, 743 were previously reported in RIC studies in plants^9,12^, whereas 392 are newly discovered here and represent novel putative RBPs. Hence, we provide a deep RBPome generated in plant tissues, which contains hundreds of putative RBPs previously unknown to bind RNA.

Flg22 treatment did not induce drastic changes in the predominant RBPs, which are detectable by silver staining analysis (**Fig. 1c**). These results are in accordance with observations from other comparative RIC studies^10,11^.

However, quantitative proteomic analyses revealed substantial alterations of the RBPome induced by flg22 (**Fig. 2a**,**b**). Out of the 1135 leaf RBPs identified, 186 (∼16% of the RBPome) displayed differential association with RNA upon immune activation (False Discovery Rate (FDR) ≤ 0.1). We refer to these here as ‘flg22-responsive RBPs’ (**Table S2**). Moreover, we identified an additional 144 RBPs (∼13% of the RBPome) with flg22-driven changes in RNA-binding with FDR ≤ 0.2, which are classified as ‘candidate flg22-responsive RBPs’ (**Table S2**). Importantly, although the candidate flg22-responsive RBPs were identified with less stringent criteria, this set contains a number of RBPs with known links to immunity such as AGO1^19^, suggesting the presence of true immune-linked RBPs. However, we recommend readers to carefully validate the responsive behaviour of these RBPs by orthogonal methods.

**Fig. 2.**
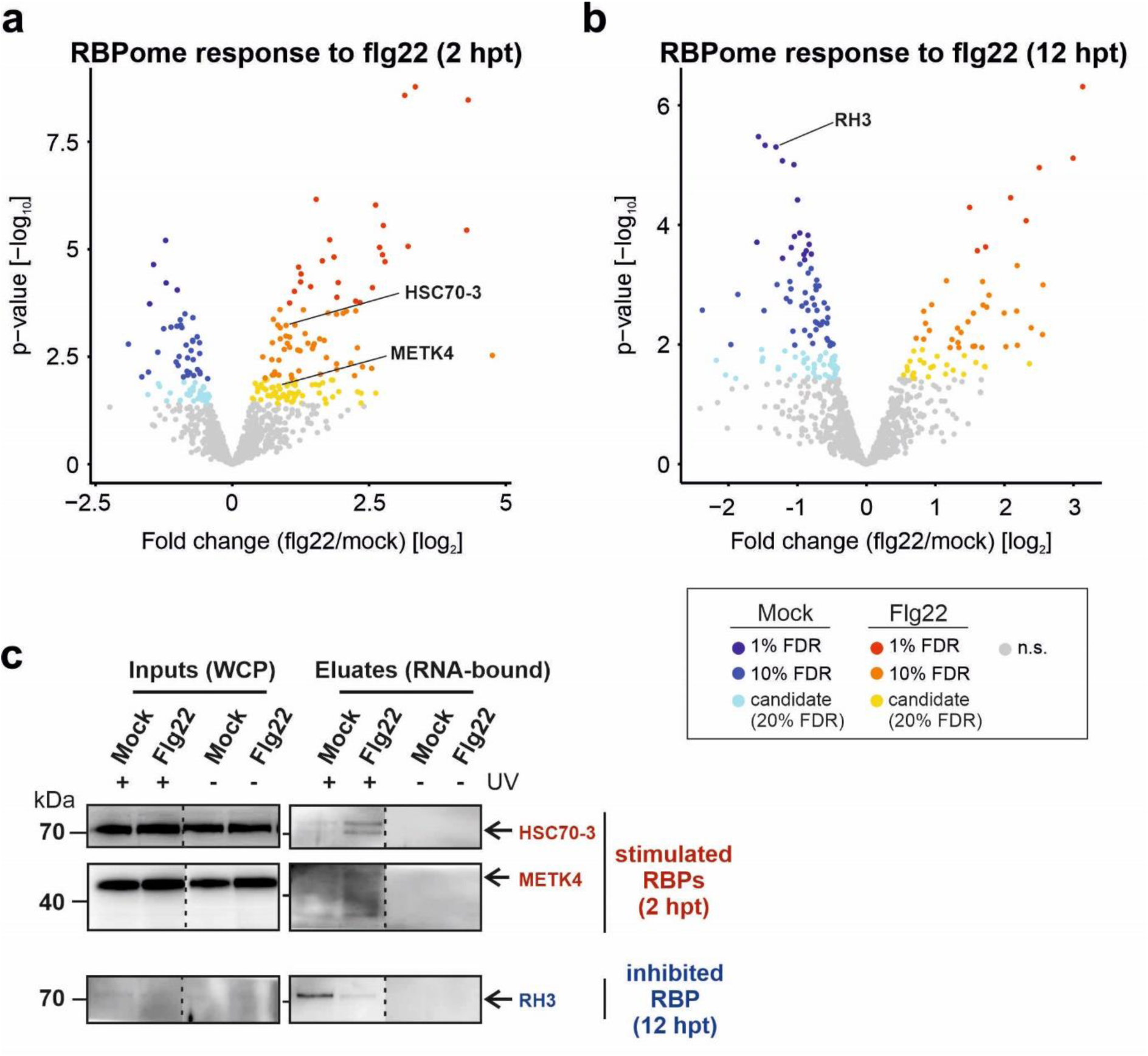
Dynamic responses of the RBPome to flg22 perception. **a-b** Volcano plots depicting the log_2_ fold change and the significance (p-value) of each protein (dots) between flg22 and mock treatment (2 hpt and 12 hpt) using data from four biological replicates. The colours of the dots indicate the significance (false discovery rate, FDR) of the protein as follows: proteins are coloured in red when FDR ≤ 0.01 and log_2_FC [flg22/mock] > 0, in orange when FDR ≤ 0.1 and log_2_FC [flg22/mock] > 0, in yellow when FDR ≤ 0.2 and log_2_FC [flg22/mock] > 0, in dark blue when FDR ≤ 0.01 and log_2_FC [flg22/mock] < 0, in blue when FDR ≤ 0.1 and log_2_FC [flg22/mock] < 0 and in light blue when FDR ≤ 0.2 and log_2_FC [flg22/mock] < 0. Red, orange and yellow proteins represent the RBPs stimulated upon flg22 perception, whereas dark blue, blue and light blue represent the RBPs inhibited upon flg22 perception. Non-significant (n.s.) proteins are coloured in grey. **c Validation of the mass spectrometry results by western blot analyses**. The leaves of mature Arabidopsis plants (5-6 weeks old) were infiltrated with either H_2_O (mock) or the immune elicitor flg22 (1 μM) and leaf tissue was harvested at 2 hpt or 12 hpt. The RNA-bound RBPs isolated by ptRIC (eluates) or the whole cell proteomes (inputs) were analysed by WB using specific antibodies against HSC70, METK4 and RH3.

Out of the 186 flg22-responsive RBPs, 112 were altered in the early response (2 hpt), 98 in the late response (12 hpt) and 24 in both (**Fig. 2a**,**b**). Moreover, out of 144 candidate RBPs, 96 were altered at 2 hpt, 71 at 12 hpt and 23 at both time points. The fact that only 24 flg22-responsive RBPs and 23 candidate RBPs were altered at both 2 and 12 hpt suggest that the plant RBPome is dynamically and temporally modulated following immune elicitation. Interestingly, 34 out of the 186 flg22-responsive proteins and 23 out of the 144 candidate proteins were detected exclusively either in mock or flg22 treated samples, suggesting that they are either induced (off/on) or repressed (on/off) in response to the elicitor. The proportion of RBPs that show differential association with RNA in response to flg22 in plants and Sindbis virus in human cells^10^ is similar, even though these experiments were done in different eukaryotic models. These results suggest that 15-30% of the eukaryotic RBPome is reactive to pathogen stimulation. Interestingly, 31 RBPs (17 of the flg22-responsive and 14 candidate RBPs) have also been shown to be differentially associated with RNA upon drought stress^16^, and may represent regulators of both biotic and abiotic stresses. These multi-stress responding proteins may be important targets for plant breeding programs to increase resistance to both biotic and abiotic stresses.

To validate the proteomic results, we performed ptRIC to isolate the RNA-bound RBPs followed by Western blotting with specific antibodies against flg22-responsive RBPs. This orthogonal analysis revealed that, in agreement with our proteomic data, both HSC70-3 and METK4 display increased association with RNA early after flg22 treatment (2 hpt), while the opposite was observed for RH3 at later times post elicitation (12 hpt) (**Fig. 2c**). Together, these results support the high quality of our proteomic data and the ability of ptRIC to capture RBP dynamics.

### Understanding RBP regulation upon flg22 elicitation

Differential RBP association with RNA can be explained by changes in RNA-binding activity, protein abundance (i.e. more protein, more binding and *vice versa*) or by changes in the target RNAs^10,11^. To test if protein abundance is a major driver of RBP responses to flg22, as occurs during *Drosophila* embryo development^11^, we analysed the inputs of the ptRIC experiment (i.e. the whole cell proteome [WCP]) by quantitative proteomics (**Fig. S2, Table S3**). While the ptRIC eluates represent the RNA-bound fraction of RBPs, the inputs provide information about the total cellular protein.

We detected and quantified in the inputs ∼71% of the RBPs identified in the eluates of ptRIC (i.e. 808 proteins), including 138 flg22-responsive proteins and 105 candidate proteins (**Fig. 3a**,**b**). Strikingly, while some (non-RBP) proteins change in abundance in response to flg22 (**Fig. S2, Table S3**), most RBPs remained unaltered, suggesting that regulation of their RNA-binding activity is the most plausible explanation for their dynamic behaviour in the ptRIC eluates (**Fig. 3a**,**b, Fig. S3**). Indeed, only 18 (out of 165) proteins displayed abundance-driven flg22-responses at the 2 hpt time point. However, at 12 hpt, 30 (out of 117) RBPs displayed abundance-driven changes in response to flg22, indicating that at later time points, changes in protein abundance gain importance in shaping the RBPome (**Fig. 3a**,**b**). The fact that remodelling of the RBPome can occur largely without changes in protein abundance has been previously observed in human cells infected with Sindbis virus^10^ and in Arabidopsis cells in response to drought stress^16^, and suggests the existence of mechanisms that alter the ability of these RBPs to engage with RNA.

**Fig. 3.**
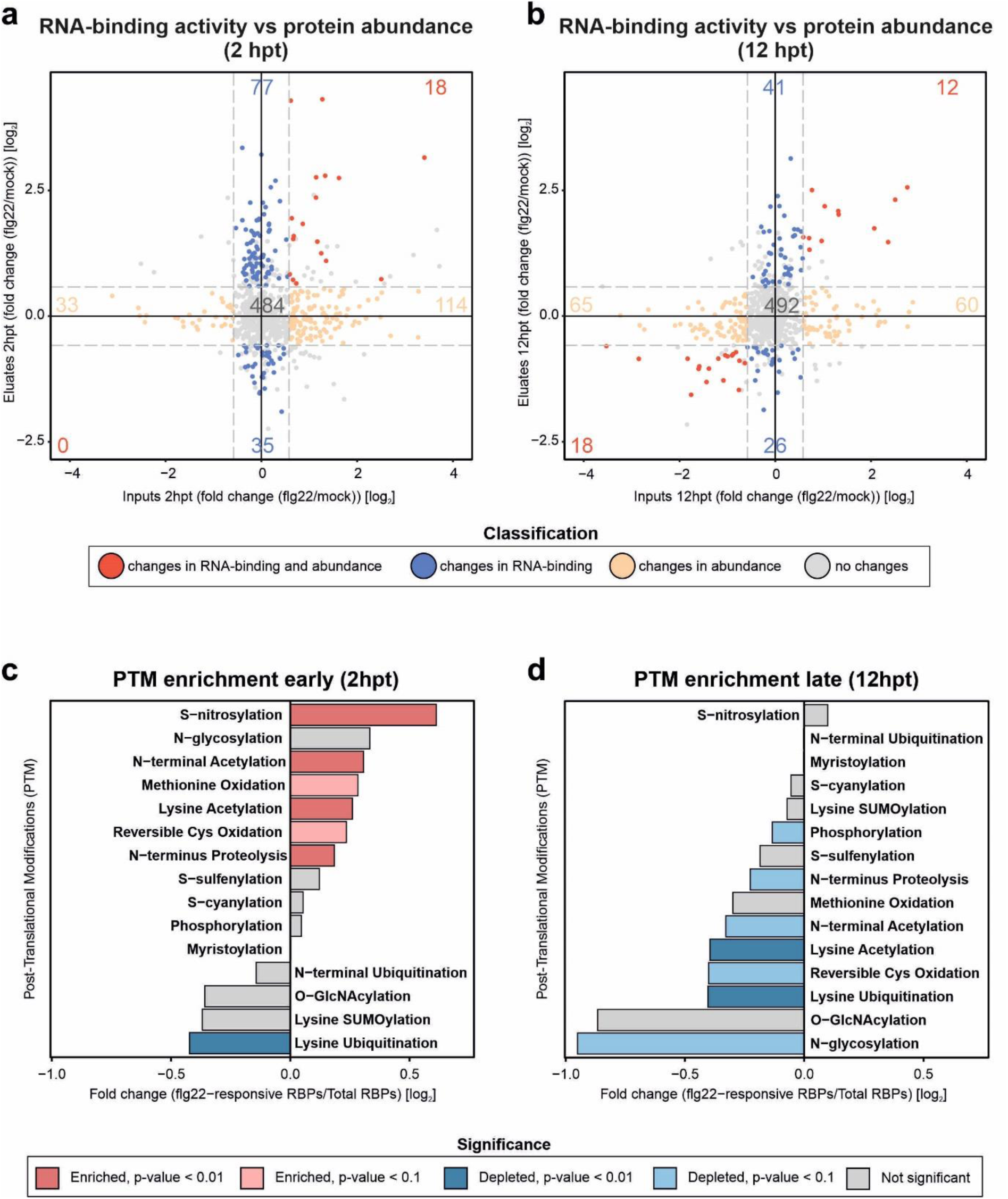
Early changes in association with RNA of the RBPome in response to flg22 do not correlate with changes in protein abundance and may be partly driven by post-translational modifications. **a-b** Scatter plots depicting the log_2_ fold change in inputs [whole cell proteome] (x-axis) and log_2_ fold change in eluates [RNA-bound RBPs] (y-axis) of each protein (dots) between flg22 and mock treatment using data from four biological replicates. Proteins are coloured according to the classification as following: proteins are coloured in red (changes in RNA-binding and abundance) when the RNA-binding activity in eluates significantly changes (FDR ≤ 0.2 and log_2_FC > 0.58 or log_2_FC < −0.58) and protein abundance changes in inputs (log_2_FC > 0.58 or log_2_FC < −0.58); proteins are coloured in blue (changes in RNA binding) when the RNA-binding activity changes (FDR ≤ 0.2 and log_2_FC > 0.58 or log_2_FC < −0.58) but the protein abundance does not change (−0.58 > log_2_FC < 0.58); proteins are coloured in yellow (changes in abundance) when the RNA-binding activity does not change (−0.58 > log_2_FC < 0.58) but the protein abundance changes in the inputs (log_2_FC > 0.58 or log_2_FC < −0.58). Non-significant proteins in eluates are coloured in grey. **c-d** Enrichment or depletion of post-translational modifications (PTMs) of flg22-responsive RBPs over total detected RBPs. Plots depicting the log_2_ fold change of the different PTM sites between the flg22-responsive RBPs and the total RBPs at 2h and 12h after flg22 treatment. Bars in dark red indicate PTMs significantly enriched with p-value ≤ 0.01, in light red significantly enriched with p-value ≤ 0.1, in dark blue significantly depleted with p-value ≤ 0.01 and in light blue significantly depleted with p-value ≤ 0.1. Non-significant PTMs are coloured in grey.

RBP activity can be regulated by post-translational modifications (PTMs) and, indeed, RNA-binding domains (RBDs) are enriched in PTM sites ^20,21^. Hence, one exciting possibility is that the signalling pathways activated by flg22 regulate RBPs through post-translational modifications (PTMs). To test this hypothesis, we analysed the distribution of PTM sites in the RBPome based on annotations from the Plant PTM viewer database^22^. Notably, PTM sites were distributed differentially in early and late flg22-responsive RBPs when compared to RBPome as a whole (**Fig. 3c**,**d**). At 2 h post elicitation, flg22-responsive RBPs were enriched in PTM sites associated with S-nitrosylation, lysine and N-terminal acetylation, N-terminus proteolysis and methionine and reversible cysteine oxidation, whereas they were depleted in lysine ubiquitination sites (**Fig. 3c**). Conversely, at 12 h post elicitation, flg22-responsive RBPs were broadly depleted in PTM sites, including those associated with lysine ubiquitination and acetylation, N-glycosylation, reversible cysteine oxidation, N-terminal acetylation, N-terminus proteolysis and phosphorylation (**Fig. 3d**). These results suggest that early changes in RBP activity may, at least partly, be controlled by PTMs, whereas PTMs may contribute less to the differential activity of RBPs at later time point.

Interestingly, enrichment in S-nitrosylation and depletion of lysine ubiquitination are common trends at both time points. S-nitrosylation consists of the reversible attachment of nitric oxide (NO) to the sulfhydryl group of cysteines. NO accumulates during plant immune responses and plays important roles in the reprogramming of the plant transcriptome^23^. For example, modification of the transcription factor SRG1 with NO via S-nitrosylation triggers an immunity-linked transcriptional programme^23^. While the importance of S-nitrosylation in transcriptional regulation has been explored, it remains unknown whether this modification also affects post-transcriptional programmes. Our data suggests that S-nitrosylation is critical for flg22-induced transcriptome reprogramming by modulating protein-RNA interactions and, consequently, post-transcriptional networks. Interestingly, 70% of the RBPs with annotated S-nitrosylation sites are stimulated upon flg22 treatment, indicating that S-nitrosylation may promote their association with RNA.

Lysine ubiquitination typically induces proteasomal-dependent protein degradation, although it can also regulate protein activity without causing proteolysis^24^. Although protein ubiquitination is important for plant immunity^24^, our data suggests that it does not play a central role in RBP regulation in response to flg22. In contrast to these results, the host cell response to human virus infection involves the activation of several E3 RNA-binding ubiquitin ligases^25^. Whether bacterial infection can trigger additional PTM regulatory layers cannot be fully recapitulated by treatment with an immune elicitor such as flg22, and thus remains to be explored with bacterial infection experiments.

Changes in the RNA-binding activity of an RBP can also be due to changes in the abundance of its target RNAs^10^. It has previously been reported that the number of differentially expressed genes dramatically increases over time after flg22 treatment^26^. Hence, we envisage that the significant changes in the abundance of thousands of transcripts at late time points could partly contribute to the changes in RNA-binding activity of late flg22-responsive RBPs.

Interestingly, 147 (2 hpt) and 125 (12 hpt) RBPs display changes in protein abundance but not in association with RNA (**Fig. 3a**,**b**). This unexpected phenomenon could be explained for example by interactions with non-poly(A) RNA, which are not captured by ptRIC, or because they are in excess over their target RNA. Either way, these results suggest that changes in abundance do not always result in matching alterations in association with RNA.

### Discovering the pathways regulated during plant immunity

UV crosslinking depends on the geometry and duration of protein-RNA interactions, as well as the nucleotide and amino acid composition at the protein-RNA interface. To determine the UV crosslinkability and the nature of interactions exerted by flg22-stimulated and inhibited RBPs, we normalised the protein intensity in ptRIC eluates (RNA-bound fraction) to that in inputs (total protein) as previously described^25,27^. Most of the RBPs for which RNA-binding activity is inhibited by flg22 treatment were enriched in ptRIC eluates over inputs (**Fig. 4a-d, Table S4**). Indeed, approximately 37% of them were exclusively detected in ptRIC eluates, which suggests that they are of low abundance in the whole cell proteome and that oligo(dT) enrichment is required to enable efficient enrichment and identification. These results suggest that flg22-inhibited RBPs establish strong and long-lived interactions with RNA, which promote high eluate/input ratios. In agreement, 65% of them have well-established links to RNA biology and are components of central RNA metabolic pathways (**Fig. 5, Fig. S4)**. This suggests that the cell down-regulates core RNA metabolic processes in response to flg22.

**Fig. 4.**
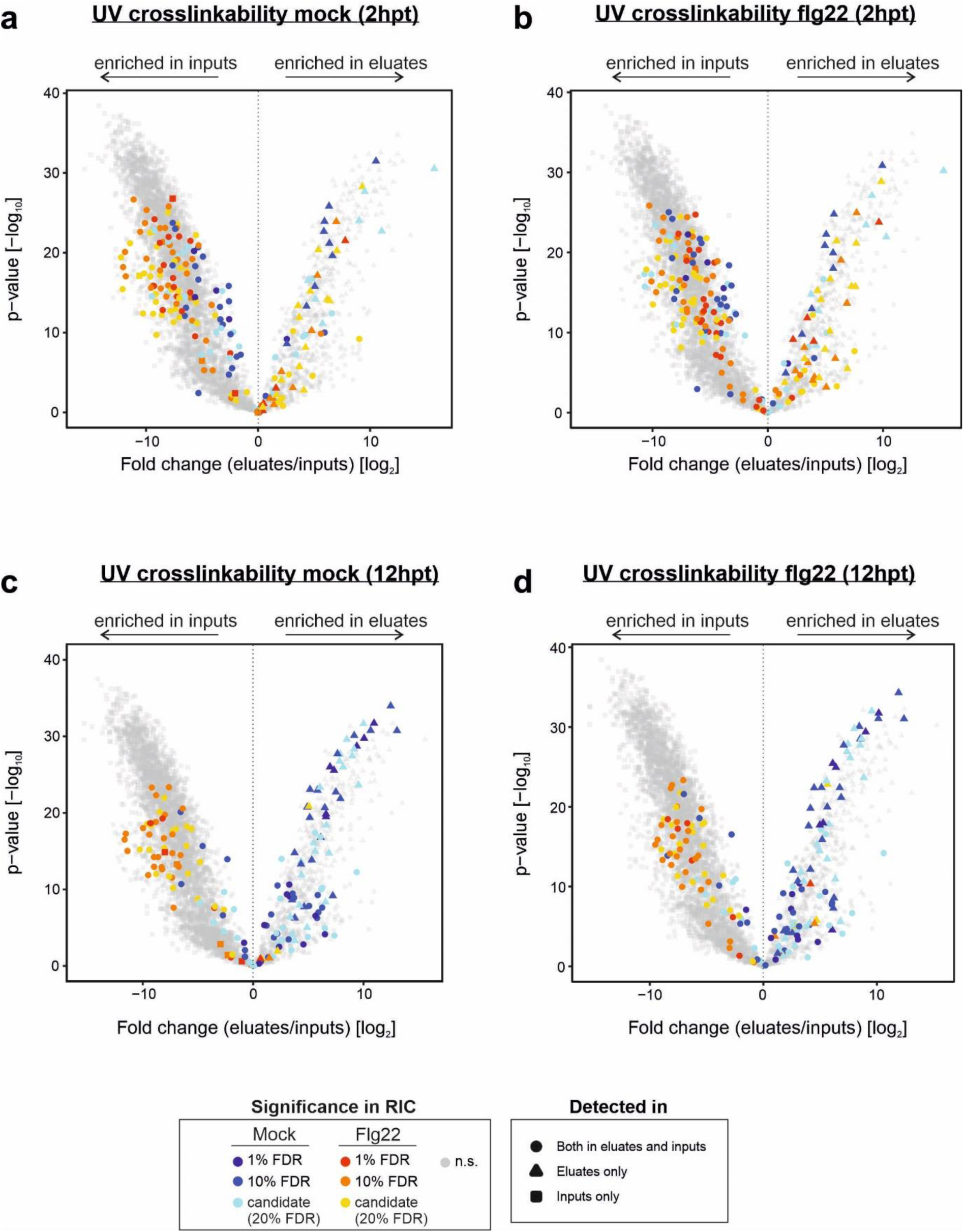
UV-crosslinkability of flg22-responsive RBPs. **a-d** Volcano plots depicting the log_2_ fold change and the significance (p-value) of each protein (dots) between ptRIC eluates (RNA-bound fraction) and inputs (total protein) at each of the different conditions (mock 2 hpt, flg22 2 hpt, mock 12 hpt and flg22 12 hpt). The colours indicate the significance (false discovery rate, FDR) of the proteins in ptRIC eluates as follows: proteins are coloured in red when FDR ≤ 0.01 and log_2_FC [flg22/mock] > 0, in orange when FDR ≤ 0.1 and log_2_FC [flg22/mock] > 0, in yellow when FDR ≤ 0.2 and log_2_FC [flg22/mock] > 0, in dark blue when FDR ≤ 0.01 and log_2_FC [flg22/mock] < 0, in blue when FDR ≤ 0.1 and log_2_FC [flg22/mock] < 0 and in light blue when FDR ≤ 0.2 and log_2_FC [flg22/mock] < 0. Red, orange and yellow proteins represent the leaf RBPs stimulated upon flg22 perception, whereas dark blue, blue and light blue represent leaf RBPs inhibited upon flg22 perception. Non-significant (n.s.) proteins are coloured in grey. The shape indicates which datasets the proteins were detected in: circles indicate proteins detected in both ptRIC eluates and inputs, triangles indicate detected only in ptRIC eluates and squares indicate detected only in the inputs. Background values were imputed for proteins not detected in any given dataset.

**Fig. 5.**
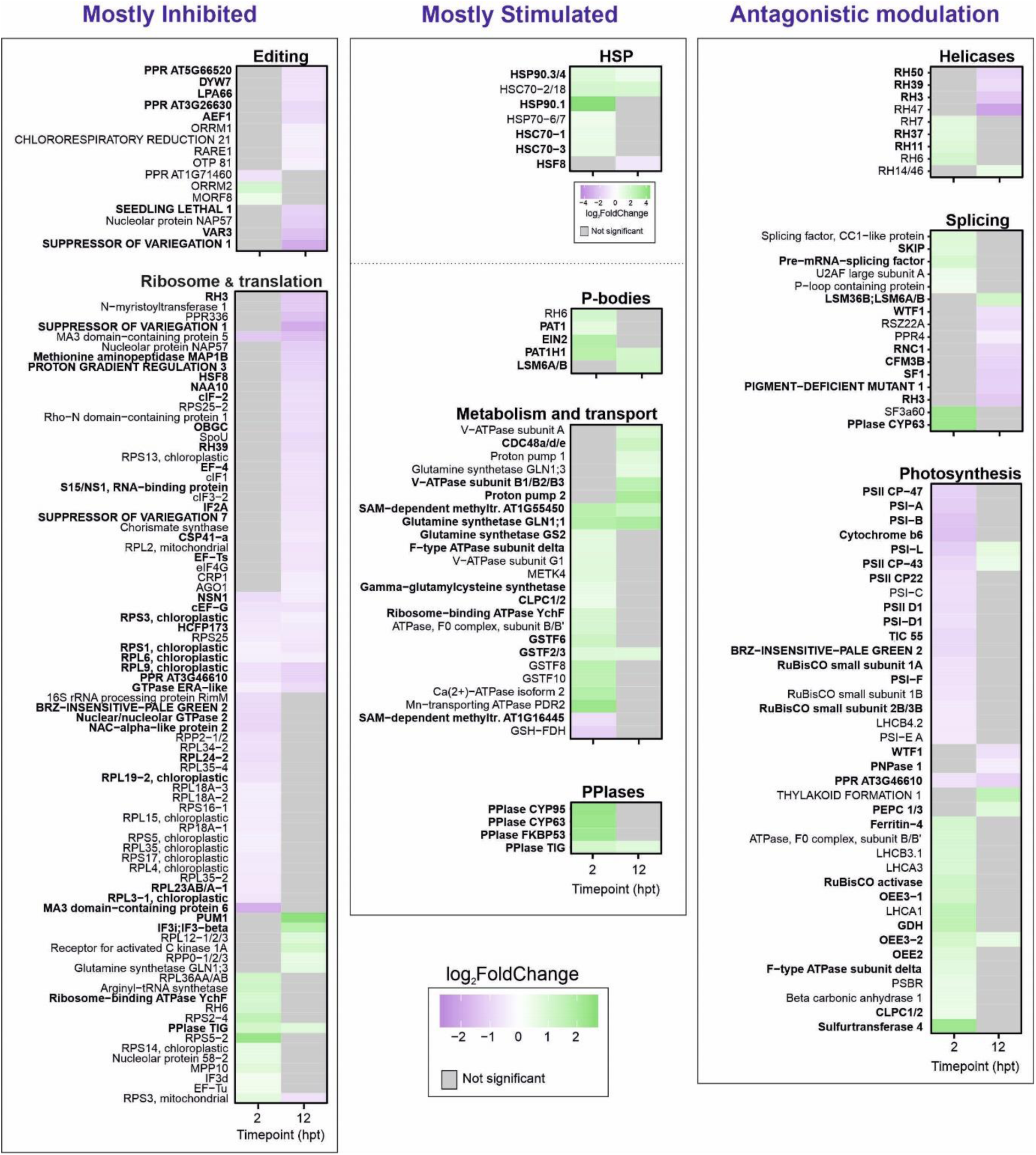
Functional categories of RBPs altered in association with RNA upon flg22 perception. Heat maps depicting the log_2_ fold change of proteins between flg22 and mock treatment (2 hpt and 12 hpt) using data from four biological replicates. Proteins are grouped into different categories based on their response to flg22 and function. Flg22-responsive RBPs detected with FDR ≤ 0.1 in any of the time points are highlighted in bold, whereas candidate flg22-responsive RBPs identified with FDR ≤ 0.2 are in regular font. Non-significant proteins are coloured in grey.

By contrast, most flg22-stimulated RBPs were detected in both ptRIC eluates and inputs and were not enriched in ptRIC eluates (**Fig. 4a-d, Table S4**). These findings suggest that either interaction with RNA is suboptimal for UV crosslinking (e.g. transitory interactions with RNA or binding to the phosphate backbone) or that only a fraction of the total protein pool actually engages with RNA upon immune activation, which is a known feature of moonlighting RBPs^28^. Accordingly, only 30% of the stimulated RBPs are associated with GO terms linked to RNA biology (**Fig. 5, Fig. S4**). Importantly, many of these flg22-stimulated RBPs have enzymatic activities (e.g. heat shock proteins [HSP], peptidyl-prolyl cis-trans isomerases [PPIases], or mono-or di-nucleotide-binding proteins [ATP/ADP, NADP/NADH]) and have been shown to bind RNA in other organisms^20,28^. Moreover, recent work has shown that these groups of proteins play important pivotal roles in response to physiological and pathological cues, including virus infection^25,28^.

To further understand the pathways altered upon flg22-induced immunity, we performed cluster analysis and analysed pathway enrichment using GO annotations and the STRING protein-protein interaction database^29^. The dynamics of flg22 response were divided into initial response and progressive response, defined by ptRIC eluate fold change from mock to early flg22 (2 hpt) and from early (2 hpt) to late flg22 (12 hpt), respectively^30^. We identified 8 clusters of proteins according to their initial and progressive responses to flg22 (**Fig. 6, Table S2**): 1) early inhibition (i.e. 2 h) followed by late recovery or stimulation (i.e. 12 h); 2) late stimulation; 3) increasing stimulation throughout the flg22 response; 4) quick inhibition reaching plateau at 2 hpt; 5) quick enhancement reaching plateau at 2 hpt; 6) increasing inhibition throughout the flg22 response; 7) late inhibition (12 hpt); and 8) rapid stimulation followed by late recovery or inhibition (i.e. 2 hpt) (**Fig. 6, Table S2**). Clusters including flg22-inhibited RBPs (i.e. 1, 4, 6 and 7) contain RBPs with defined roles in RNA metabolism, including RNA editing and translation. Additionally, cluster 1 includes several proteins involved in photosynthesis and cluster 7, high numbers of PPR-containing proteins.

**Fig. 6.**
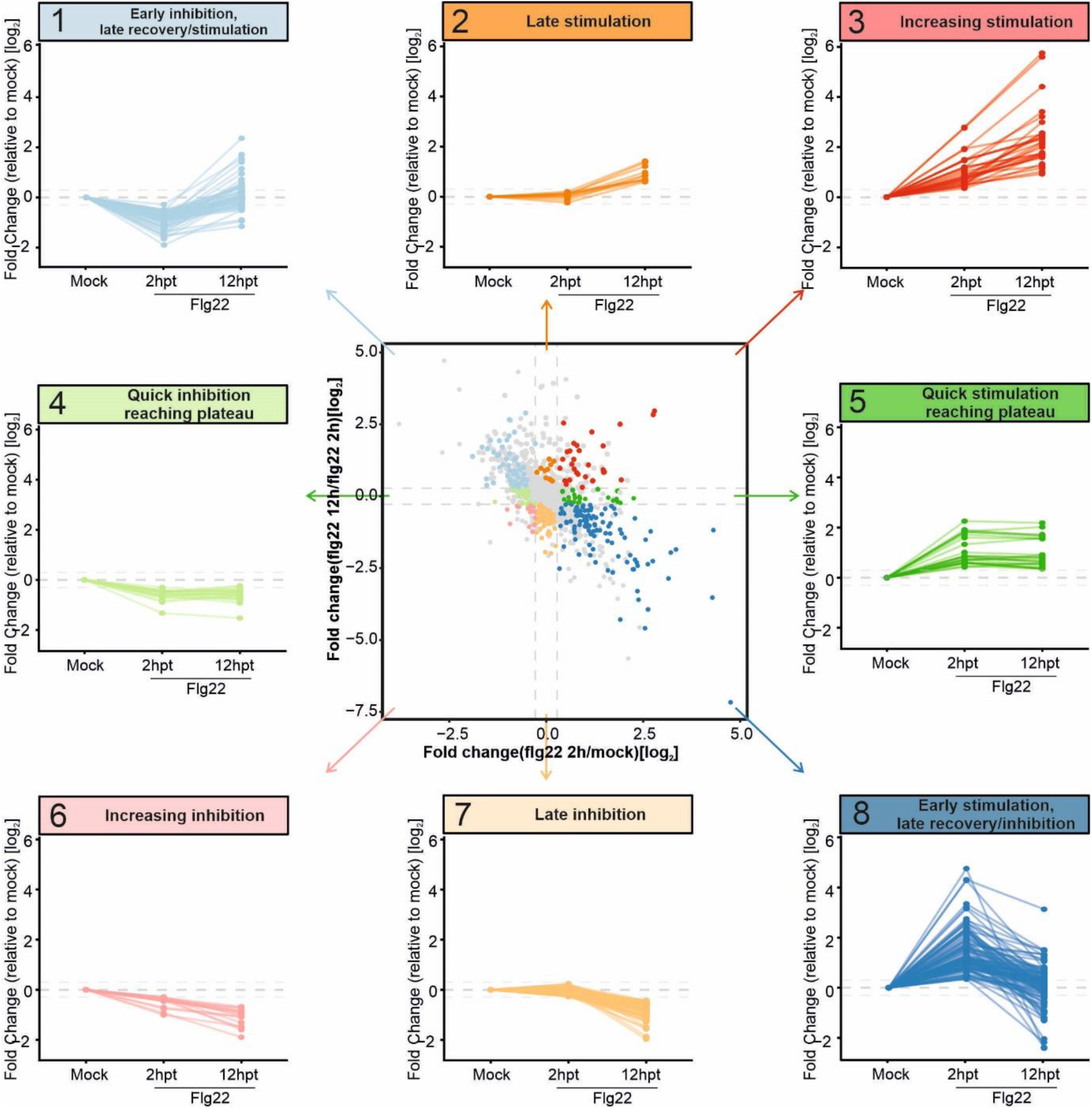
Dynamics of RNA-binding activity in flg22 responsive RBPs. The flg22-responsive RBPs were divided into 8 clusters based on their initial response (log_2_ fold change of flg22 2h/mock) and progressive response (log_2_ fold change of flg22 12h/flg22 2h) to flg22^30^: 1) early inhibition (i.e. 2h) followed by late recovery or enhancement (i.e. 12h); 2) rapid stimulation followed by late recovery or inhibition (i.e. 2h); 3) quick inhibition reaching plateau at 2hpt; 4) quick enhancement reaching plateau at 2hpt; 5) Increasing inhibition throughout flg22 response; 6) increasing stimulation throughout flg22 response; 7) late inhibition (12h); and 8) late stimulation.

Conversely, the clusters involving stimulated RBPs (i.e. 2, 3, 5 and 8) were enriched in stress response proteins (including biotic stress) and moonlighting enzymes. Taken together, these data suggest the RBP responses to flg22 are clustered based on protein function. How this coordinated regulation contributes to immunity deserves detailed characterisation in the future and is further discussed below.

### Flg22 treatment inhibits RBPs involved in editing and protein synthesis

In plants, plastid and mitochondrial transcripts can be edited post-transcriptionally, which often results in amino acid substitutions in protein sequences. Multiple RBPs involved in RNA editing, mostly in the chloroplast, were inhibited after 12 h of flg22 treatment (**Fig. 5, Table S2**). These included 11 proteins involved in C-to-U RNA editing, 2 pseudouridine synthases and 3 proteins annotated by ‘RNA modification’ but largely uncharacterised. Most of these proteins (12 out of 16) belong to cluster 7 (**Fig. 6**). For example, we found inhibition of the ORRM1 (Organelle RRM protein 1), which controls 62% of the chloroplastic C-to-U RNA editing sites^31^, and of VAR3 (VARIEGATED 3), which is a zinc-finger motif containing protein that controls a large number of chloroplastic RNA editing sites^32^. These results agree well with earlier studies showing that pathogen challenge inhibits chloroplastic RNA editing in plants as a response to increase pathogen resistance^33^.

Plants can adapt to environmental changes by regulating mRNA translation, which allows rapid control of protein abundance^34^. Interestingly, flg22 induces changes in RNA-binding activity of 8 eukaryotic initiation factors (IFs) and 4 elongation factors (EFs), 32 ribosomal proteins and several proteins associated with the ribosome, most of which are inhibited (**Fig. 5, Table S2**). Moreover, recent studies using ribosome profiling have reported that translation can be selectively regulated during plant immune responses^34–36^. Hence, our data provides further insights into translation regulation during plant immunity, showing that the RNA-binding activity of multiple components of the translation machinery, including ribosomal proteins, is altered in response to flg22. A similar global translational slowdown has been suggested to play important roles as an antiviral strategy^37^ and to occur during responses to abiotic stresses such as hypoxia^38^, heat^39^ or drought^40^. We hypothesise that this phenomenon may play a role in or be a consequence of the described switch from translation of growth-related mRNAs to those involved in rapid adaptive responses^36^.

Nascent peptides can be modified co-translationally at the *N*-terminal, affecting for example the protein half-life and folding characteristics. In eukaryotes two major protein modifications are *N*-terminal Met excision carried out by Met aminopeptidases (MAP) and *N*-α-terminal acetylation catalysed by *N*-terminal acetyltransferases (NATs)^41,42^. Our data indicates inhibition of the chloroplastic Met aminopeptidase MAP1B, and the cytoplasmic NAA10, which is the catalytic subunit of *N*-terminal acetyltransferase A (NATA). Both MAP1B and NAA10 were previously found to interact with RNA in plants^12^. Interestingly, NAA10 is involved in defence against pathogens by regulating acetylation of the first Met of the disease resistance proteins SNC1 and RPM1, thus targeting these proteins for degradation^43^. Hence, plants may actively inhibit the association of NAA10 with RNA as part of flg22-triggered immunity to suppress degradation of proteins important for plant immune responses. Another common protein modification is *N*-myristoylation (MYR) by *N*-myristoyltransferases (NMTs), which generally acts to target proteins to membranes. We identified flg22-driven inhibition of the cytoplasmic protein NMT1, which *N*-myristoylates different immune receptors (NLRs)^44^. It has been shown that NMT1 interacts with ribosomes and this interaction is mediated by RNA^45^. The role of the differential interaction between nascent peptide modifying enzymes and RNA in the context of plant immune response deserves further characterisation.

### Stimulation of RBPs involved in processing bodies by flg22

P-bodies are dynamic cytoplasmic assemblies that comprise translationally inactive mRNAs and proteins involved in mRNA turnover and translational repression^46^. It is known that during stress conditions some mRNAs can be re-localised to P-bodies^46,47^ and recent studies have demonstrated that P-bodies greatly contribute to immune responses by rapidly degrading or storing some target mRNAs^4,48^. Importantly, flg22 treatment stimulated the RNA-binding activity of several proteins associated with processing bodies (P-bodies), including PAT1, PAT1H1, LSM6A/B, EIN2, and RH6 (**Fig. 5, Table S2**).

PAT1 is one of the three Arabidopsis homologs of the yeast PAT1 and functions as an enhancer of decapping by binding the 3’ end of deadenylated mRNAs together with LSM1-7 and promoting decapping of specific transcripts^49,50^. In addition, PAT1 plays a role in translational inhibition and P-body formation during flg22-elicited immune responses^50^. We observed that PAT1, and another Arabidopsis PAT1 homolog PAT1H1, associate with RNA to a greater extent upon flg22 treatment. Interestingly, the human PATL1 is also stimulated after virus infection correlating with a pervasive remodelling of the host cell transcriptome^10^, which suggests that this is a common theme in pathogen response across eukaryotes. We also detected stimulation of the RNA helicase RH6, which was recently discovered to regulate plant immunity through the control of decapping-dependant mRNA decay^51^. Taken together, our results suggest that critical P-bodies components react functionally to flg22 treatment, suggesting a potential role in rewiring the cell transcriptome.

### Several enzymes moonlight as RBPs in response to flg22 treatment

In recent years, it has become apparent that proteins lacking classical RNA-binding architectures can interact with RNA through unconventional RNA-binding modes^9,28^. Indeed, a recently developed method to profile RBDs at a proteome-wide scale has shown that RNA-binding surfaces often co-occur with enzymatic activities and protein-protein interaction domains^20^. These findings agree well with earlier discoveries showing that several metabolic enzymes moonlight as RBPs, often under particular cellular states such as stresses^28^. For example, during iron deficiency or oxidative stress the animal aconitase/IRP1 becomes an RBP that regulates mRNAs related to iron homeostasis^9^. Here, we show that several proteins harbouring enzymatic activity display an enhanced RNA-binding activity after flg22 treatment in plant leaves (**Fig. 5, Table S2**). This striking result suggests that moonlighting RBPs can also be a common theme in plant physiology.

There are two well-established functional routes for moonlighting RBPs. First, they can regulate gene expression at the post-transcriptional level, as occurs with IRP1^9^. Alternatively, the interaction with RNA can allosterically regulate protein function^9^ as occurs with the autophagy factor p62^52^ or the E3 ubiquitin ligase TRIM25^53^. Below, we discuss several moonlighting RBPs that are stimulated by flg22. Whether they are post-translational regulators or are regulated by RNA deserves further characterisation.

PPIases are molecular chaperones with important roles in protein folding and regulation by mediating the *cis* to *trans* transition of peptidylprolyl bonds. Recently, PPIases have been classified as RBPs from yeast to human^9,20^ and have been shown to play critical roles in RNA metabolism^54^. Moreover, for some PPIases their RNA-interacting surfaces have been described^20^ or contain well characterised RNA-binding domains (RRM), which reinforces the intimate connection between PPIases and RNA. However, whether PPIases display RNA-binding activity in plants remains poorly understood. Our data shows that at least 7 plant PPIases interact with RNA in leaves (**Table S1**) and the RNA-binding activity of four of them (FKBP53, TIG, CYP63 and CYP95) is stimulated by flg22 (**Fig. 5, Table S2**). This agrees with previous findings in virus-infected human cells, where PPIA’s RNA-binding activity is stimulated and regulates viral gene expression^10^. Moreover, an effector from the cyst nematode *Heterodera schachtii* targets the flg22-stimulated PPIase FKBP53^55^, suggesting its importance in immunity (**Fig. 5**). These observations highlight the importance of PPIases for plant immunity, although their links to RNA biology remain to be elucidated.

HSPs are molecular chaperones that facilitate protein folding. Surprisingly, recent studies have shown that multiple members of the HSP70 and HSP90 families interact with RNA in plants^12^ and humans^7,9^, and for some human HSP70/90 proteins their RNA-binding surfaces are known^20^. Here, we identified 11 HSPs that interacted with RNA in plant leaves (**Table S1**), 6 of which were stimulated upon flg22 treatment. Stimulated HSPs included members of the HSP90 and HSP/HSC70 families (**Fig. 5, Table S2**). This activation may reflect a role for HSPs in managing protein unfolding derived from immunological responses. Similarly, several human HSPs display differential RNA-binding activity upon RNA virus infection, and HSP90AB1 was shown to be crucial to support viral gene expression^10^. In plants, HSP70 and HSP90 have been associated with immunity^56^ and targeting of HSP90 by a *Pseudomonas syringae* effector disrupts the plant immune response^57^. Taken together, these findings strongly support the participation of HSPs as regulators of immunity, although the role of their RNA-binding activity remains to be solved. Increased interaction of HSPs with RNA during immune responses may cause or arise from the association of HSPs with ribonucleoprotein complexes (RNPs). RBPs are highly disordered proteins and many of them have the propensity to condense into RNA granules^58^. Therefore, the association of HSPs with RNPs may help to prevent protein aggregation and cellular stress as previously suggested^20^.

Interestingly, our data revealed the activation of several proteins that moonlight as RBPs in response to flg22. Multiple glutathione S-transferases (GSTs) displayed enhanced interaction with RNA after flg22 perception (**Fig. 5, Table S2**). Glutathione (GSH) plays an important role in immune signalling and some studies have established links between GSH and RNA regulation^59^. For example, GSH modulates the synthesis of the key immune signalling molecule ethylene by regulating the mRNA stability of the biosynthetic enzyme ACO1^59^. Moreover, several ATPases with RNA-binding activity were stimulated upon flg22 treatment, some of which contribute to the acidification of the apoplast in response to pathogen attack (**Fig. 5, Table S2**). Few ATPases have been reported to bind poly(U)^60^, however, the role of ATPases in RNA metabolism remains unknown. In addition, several proteins involved in glutamine metabolism displayed increased RNA-binding activity upon flg22 treatment, including the cytosolic glutamine synthetases GLN1-1 and GLN1-3, and the chloroplastic GLN2 (**Fig. 5, Table S2**), suggesting that protein-RNA interactions could affect the role and regulation of these enzymes in metabolism and immunity.

### Modulation of RNA helicases, RNA splicing factors and photosynthesis-related proteins in response to flg22

While the protein groups described above showed a unified response type to flg22, other groups and pathways contained proteins that reacted differentially to flg22 treatment (**Fig. 5, Fig. S3, Table S2**). These RBPs included canonical RBPs as well as unconventional RBPs with previously unknown roles in RNA biology.

RNA helicases are crucial at many steps of RNA metabolism since they catalyse the unwinding of RNA secondary structure. Hence, changes in their activity are expected to affect the remodelling capacity of RNPs^61^. We identified 9 RNA helicases (around 15% of Arabidopsis helicases) as differentially regulated upon immune elicitation, five of which were stimulated and four inhibited (**Fig. 5, Table S2**). Most of the inhibited RNA helicases had a chloroplastic localisation, whereas the stimulated RNA helicases were nuclear or cytosolic. In agreement with our results, 16 RNA helicases displayed altered RNA-binding activity in SINV-infected human cells^10^. Our data show that the chloroplastic DEAD-box RNA helicase RH50 is inhibited 12 h after flg22 perception. Interestingly, the RH50 rice ortholog, OsBIRH1, is known to play important roles in biotic and oxidative stress^62^, suggesting that the role of RH50 in immunity might be conserved from monocots to dicots.

RNA splicing and its regulation are of paramount importance for the adaptive response to different environmental, physiological and pathological challenges^63^. Here we identified 16 flg22-responsive RBPs linked to splicing (**Fig. 5, Table 22**). Interestingly, nuclear splicing factors are stimulated early upon flg22 treatment. Conversely, chloroplastic splicing factors are inhibited at 12 h post treatment. Four flg22-responsive splicing factors are annotated to be involved in stress responses (SKIP, SF1, RH3 and U2AF large subunit A). One example is the evolutionarily conserved SKI-INTERACTING PROTEIN (SKIP) that functions as both a transcription and splicing factor and regulates alternative splicing of genes related to tolerance to abiotic stresses^64^. Differential regulation of splicing factors has also been observed in human cells infected with Sindbis virus^10^ and Arabidopsis cell cultures upon drought stress^16^. Thus, splicing emerges as a tightly regulated process in response to pathogens and stresses.

We recently reported that photosynthesis-related proteins interact with poly(A) RNA in plants leaves^12^. Interestingly, our present study reveals that many proteins from the photosynthetic apparatus display differential association with RNA at 2 h after flg22 treatment (**Fig. 5, Table S2**). Nineteen components of the photosynthesis pathway exhibited reduced RNA-binding activity at 2 hpt, including proteins within PSII, PSI, rubisco subunits and cytochrome b6. By contrast, the opposite was observed for 15 proteins within the oxygen-evolving complex, light-harvesting complex, and F-ATPase (**Fig. 5, Table S2)**. Remarkably, the RNA-binding activities of most of these proteins returned to normal at 12 hpt, suggesting that these moonlighting RBPs are regulated dynamically throughout the flg22-induced response. While representing an exciting observation, the biological meaning of this dynamic behaviour deserves further investigation. It is worth noting that Rubisco large subunit (LSU) and cytochrome f have been reported to regulate their own RNAs in specific conditions^65,66^, and it is possible that other chloroplastic proteins engage in similar regulatory feed-back loops, in analogy with the human thymidine synthase^67^.

### Flg22-responsive RBPs are important for plant immunity

To test if the flg22-responsive RBPs contribute to plant immunity, we examined whether Arabidopsis knock out mutant lines for a subset of these proteins showed altered disease resistance. We selected 19 flg22-responsive RBPs based on the availability of mutants (**Table S5**), 15 from the high-confidence (10% FDR) and 4 from the candidate (20% FDR) responsive RBP group. Two of the selected RBPs were identified as protein groups instead of unique proteins due to their sequence similarities (HSP90-3/4 and HSP70-2/18), and therefore, mutant lines of all the RBPs within the protein group were screened. None of the mutants had a severe developmental phenotype (**Fig. S5c**,**f**,**h**). The mutants were screened for their susceptibility to the model pathogen *P. syringae* pv. tomato DC3000 (*Pst*DC3000) and its mutant hrpA^-^. *Pst*DC3000 causes disease in Arabidopsis whereas the non-pathogenic hrpA^-^ mutant elicits PTI^68^.

Out of the 21 RBPs screened, 11 had a phenotype of altered resistance or susceptibility to at least one of the two bacterial strains tested (**Table S5**). These included 3 RBPs whose mutants displayed higher resistance (RH11, METK4 and TSS), and 8 RBPs with higher susceptibility (EIN2, EFG, ClpD, HSP90-3, HSC70-1, RRM-containing protein, DHBP synthase RibB-like alpha/beta domain-containing protein and RNA polymerase II degradation factor-like protein). Interestingly, 56% of the screened mutants for the high-confidence (10% FDR) and 40% for the candidate group (20% FDR) caused alterations in the disease phenotype (**Table S5**). These results indicate that even if candidate flg22-responsive RBPs are identified with 20% FDR, they include *bona fide* immunity-linked proteins that may deserve further investigation after appropriate orthogonal validation. We selected 3 mutants with altered disease resistance phenotypes for further characterisation. These were selected as a canonical RBP (RH11), a novel RBP (METK4) and a recently described RBP with an unusual mode of RNA binding (EIN2).

The DEAD-box ATP-dependent RNA helicase RH11 (RH11) displayed increased association with RNA at 2 h after flg22 perception with no changes in protein abundance (**Table S2**). Interestingly, RH11 has an annotated S-nitrosylation site^22^, which is a common feature for RNA helicases^69^, and which could be a possible regulatory mechanism. Although RH11 remains poorly characterized, recent studies have linked RH11 to various processes, including mRNA decay, splicing or ribosome biogenesis^70–72^. To confirm the bacterial infection phenotype, we tested two independent *rh11* mutant lines, revealing that loss of RH11 activity increases resistance to *Pst*DC3000 strain but not to hrpA^-^ (**Fig. 7a-b, Fig. S5a-c**). This suggests that RH11 is involved in a process linked to effector-mediated susceptibility. Moreover, *rh11* mutants had an increased ROS burst (**Fig. 7c**), indicating that RH11 may control the RNA metabolism of genes associated with oxidative burst.

**Fig. 7.**
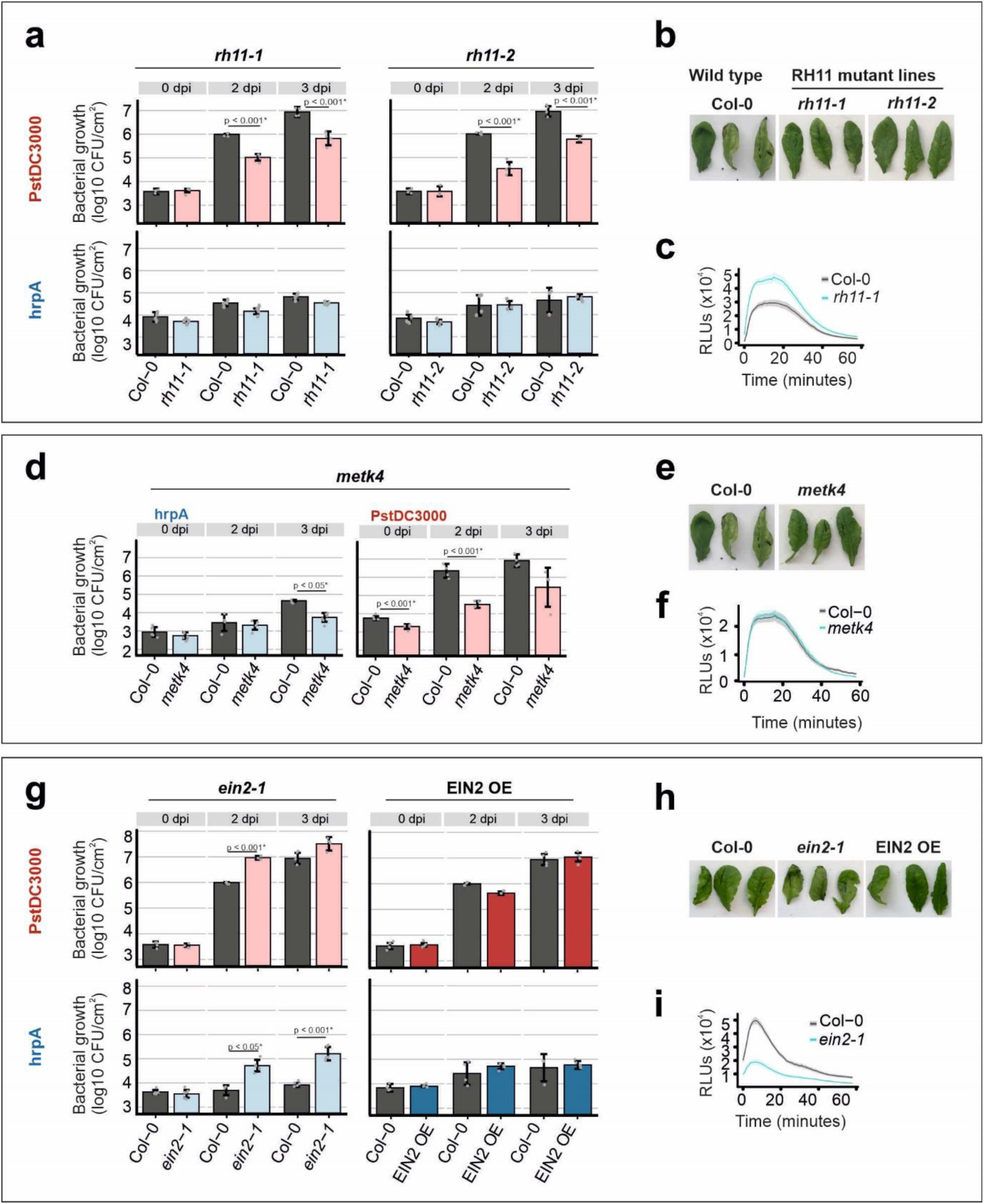
Multiple flg22-responsive RBPs are involved in immunity in Arabidopsis. **a-c** *rh11* mutants are more resistant to the disease-causing strain *Pst*DC3000 but not to its *hrpA-* mutant and display an increased ROS burst. **d-f** The *metk4* mutant is more resistant to the disease-causing strain *Pst*DC3000 and the *hrpA*-mutant. **g-i**) The *ein2-1* mutant is more susceptible to the disease-causing strain *Pst*DC3000 and the *hrpA*-mutant, and displays a decreased ROS burst, while the EIN2 overexpression line (EIN2OE) does not show any change in resistance. **a, d, g** Mutant lines, overexpression lines and Col-0 plants were infiltrated with *Pst*DC3000 and its *hrpA-*mutant at an inoculum density of 5×10^5^ cells/ml and the bacterial density analysed at 0, 2 and 3 days after inoculation (dpi). Error bars indicate mean ± sd of n = 4. ANOVA with Tukey HSD was used to calculate the p-value. The experiment was repeated at least twice with similar results. **b, e, h** Infection phenotype of mutant lines and overexpression lines at 3 dpi. **c, f, i** The ROS burst in Col-0 and mutant lines after elicitation with flg22 (500 nM). Error bars indicate mean ± SEM of n = 12. The experiment was repeated multiple times with similar results.

S-adenosylmethionine synthase 4 (METK4) association with RNA was increased at 2 h after flg22 treatment with no changes in protein abundance (**Table S2**). S-adenosylmethionine synthases catalyse the synthesis of S-adenosylmethionine (SAM), which is important both in methionine metabolism and as a universal methyl group donor molecule for a broad range of biological reactions^73^. In addition, SAM-dependent methyltransferases can transfer the methyl group to DNA, RNA, proteins or other metabolites^73^. Interestingly the RNA-binding activity of two S-adenosyl-L-methionine-dependent methyltransferases (AT1G16445 and AT1G55450) was altered upon flg22 perception, AT1G16445 being inhibited whereas AT1G55450 was stimulated (**Table S2**). METK4 has been shown to be an important player in DNA and histone methylation^74^ and interacts with TAD3, a tRNA adenosine deaminase involved in tRNA editing^75^. Although METK4 does not possess any known RBD, we previously reported that METK4 can bind RNA in Arabidopsis leaves^12^ and we have validated this finding with independent ptRIC experiments followed by proteomics or WB (**Fig. 2c**). Contrarily to what was previously reported for CRISPR/Cag9 generated *metk4* mutants^74^, our *metk4* mutant line (Salk_052289C) was viable. Strikingly, *metk4* mutant plants were more resistant to both *Pst*DC3000 and hrpA^-^ (**Fig. 7d**,**e, Fig. S5d-f**). Bacterial resistance was not linked to the ROS burst, since the ROS levels were comparable to those of the wild type plants (**Fig. 7f**). Our results indicate that METK4 is involved in plant immunity.

EIN2 was stimulated early after flg22 perception (2 hpt) (**Table S2**). Unfortunately, EIN2 was not detected in the whole cell proteome, suggesting that it is not abundant. EIN2 is a component of the ethylene signalling pathway that is conserved through evolution^76^ and has been extensively linked to responses to many environmental cues in different species, including responses to biotic stress^77,78^. EIN2 regulates FLS2, the receptor responsible for sensing flg22, and *ein2* mutants accumulate lower levels of FLS2 mRNA and protein^79,80^. Moreover, *ein2* mutants have been described to have reduced immune responses and to be more susceptible to *Pst*DC3000, especially at early time points^79,80^, suggesting that EIN2 is important in the early stages of plant immunity. However, there is controversy with respect to the *ein2* mutant phenotype since other studies have reported opposite effects in infection^81^. In our experimental set up, e*in2-1* mutants were more susceptible to infection by *Pst*DC3000, especially at the early time point (2 days post infection), and to hrpA^-^ at both early and late time points (**Fig. 7g-h, Fig. S5g**,**h**). Overexpression of EIN2 (EIN2OE) did not alter resistance to *Pst*DC3000 or hrpA^-^ (**Fig. 7g-h**). This supports the findings by Mersmann and colleagues and Tintor and colleagues^79,80^. Moreover, loss of EIN2 in *ein2-1* mutant caused a consistently reduced ROS burst in response to flg22 (**Fig. 7i**).

EIN2 contains an ER-localised transmembrane domain and a cytoplasmic domain. Upon ethylene perception a portion of the cytoplasmic domain, the CEND, is cleaved and two modes of action have been described for the EIN2 CEND (**Fig. S6**). Firstly, the CEND can re-localise to the nucleus and stabilize two transcription factors (EIN3/EIL1) that positively regulate ethylene responses^82–84^. Secondly, the CEND can remain cytoplasmic and bind the 3’ UTR of mRNAs that negatively regulate ethylene responses (EBF1/2). The CEND re-localises to the P-bodies, together with the NMD machinery, and promotes translational repression of those mRNAs^85,86^. EBF1/2 promotes degradation of EIN3/EIL1 transcription factors. Therefore, by promoting translational repression of EBF1/2, EIN2 promotes the expression of ethylene responsive genes.

To profile the responses of EIN2 upon flg22 perception, we mapped the peptides identified by proteomics and their intensities to EIN2 in the 4 different conditions (mock 2 hpt, flg22 2 hpt, mock 12 hpt and flg22 12 hpt). As expected from ptRIC eluate analyses, we identified more peptides and, generally, with higher intensity in plants treated with flg22 than with H_2_O (**Fig. 8a**). Moreover, intensity was back to mock levels at 12 hpt, suggesting that the EIN2 response to flg22 is rapid and dynamically regulated. These results align well with previous findings suggesting that EIN2 is involved in the early preinvasive immunity response^80^. Interestingly, the peptides detected were not restricted to the CEND, but mapped to the whole cytoplasmic domain (**Fig. 8a**). This suggests that either the cleavage site is not at position 645 as previously described^83^, or that an alternative cleavage site that includes the complete cytoplasmic domain exists and that it is also able to interact with RNA.

**Fig. 8.**
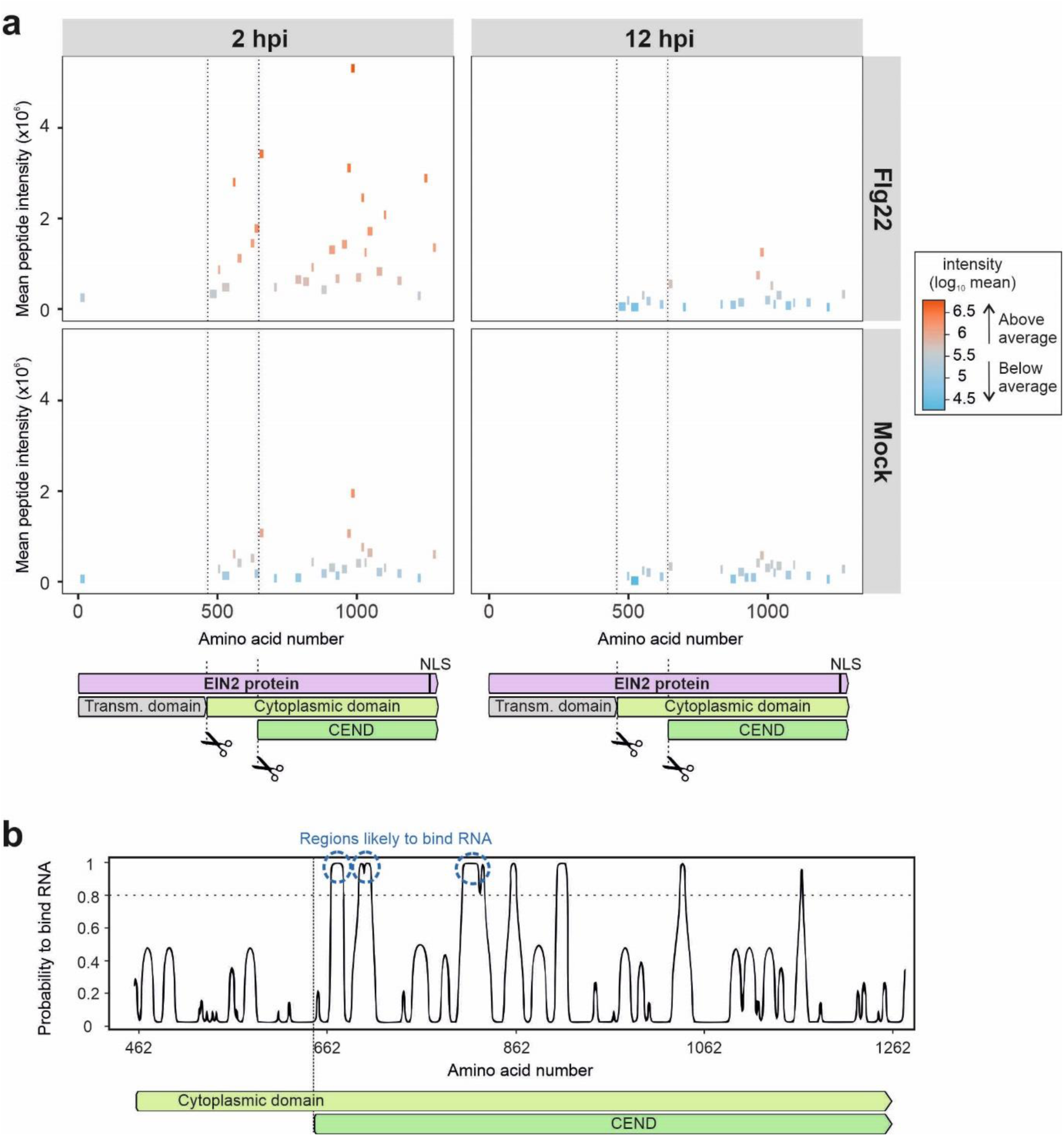
EIN2 is stimulated at early time points following flg22 perception and is cleaved at the C-terminal domain. **a** Mapping of the peptides identified by MS to the EIN2 sequence and mean peptide intensity using data from four biological replicates. Colours indicate the log_2_ mean intensity of the peptides. Dotted lines indicate the cleavage sites of EIN2: a known cleavage site that yields the CEND, and a newly identified putative alternative cleavage site at the N-terminal part of the cytoplasmic domain. **b** Prediction of the RNA-binding regions of the EIN2 cytoplasmic domain. Abbreviations: NLS, nuclear localization signal.

Importantly, the intensity of all peptides mapping to the EIN2 cytoplasmic domain in the eluates of ptRIC is similar, even upstream of the N-terminal boundary of the CEND domain (**Fig. 8a**). These results suggest that the whole cytoplasmic domain, instead of the CEND domain alone, engages with RNA in response to flg22. While our proteomic data cannot fully rule out that the full-length protein interacts with RNA, the substantially lower intensity of the single peptide mapping to the N-terminal transmembrane region indicates that an RNA-binding protein product compressing the whole cytoplasmic domain exists. Whether this is derived from proteolytic cleavage similarly to the CEND or to a different regulatory mechanism such as alternative splicing, deserves future in detail characterisation. Using an algorithm trained with RBDmap data^87^, we predicted that the cytoplasmic domain of EIN2 harbours 3 putative RNA-binding sites, and that all these regions fall within the CEND (**Fig. 8b**).

## OUTLOOK

Using ptRIC, we discovered 186 RBPs with altered RNA-binding activity upon treatment with flg22. Moreover, we identified 144 additional candidate RBPs to be potentially regulated by flg22. Flg22-responsive RBPs include proteins involved in virtually every step of RNA metabolism, from synthesis to decay, as well as unconventional RBPs previously unknown to bind RNA (**Fig. 9**). However, for most of these RBPs, their precise role in RNA metabolism and/or immunity is unknown. To understand their mechanism of action, it is now critical to test their effects in bacterial infection and to identify their target RNAs. Several crosslinking, immunoprecipitation (CLIP) and sequencing methods have been established in the last few years. Applying these techniques to flg22-responsive RBPs in flg22-treated leaves will allow identification of the RNA network modulated during immune responses^88^, and contribute to understanding of how RBPs control or are controlled by RNA in response to pathogen perception.

**Fig. 9.**
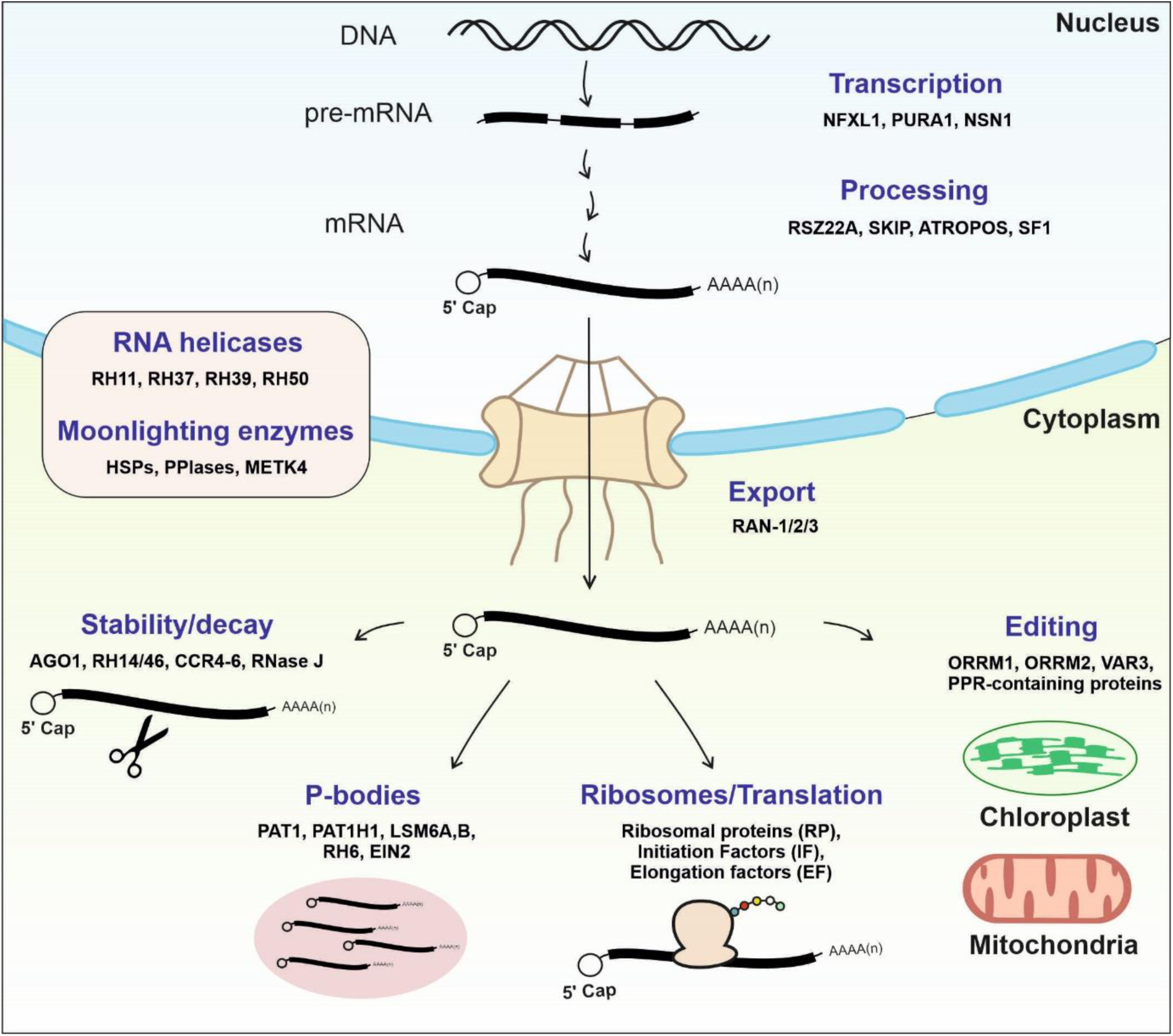
Flg22-responsive RNA-binding proteins. Flg22-responsive RBPs include proteins that play important roles in all the steps of RNA metabolism and localization including transcription, RNA processing, mRNA export, RNA stability/decay, localization to P-bodies (PBs) or stress granules (SGs), RNA editing (mainly in organelles), ribosomes and translation. RNA helicases and enzymes moonlighting as RBPs are involved in multiple processes in different cellular compartments.

Plants are constantly challenged by pathogens and their immune system responds to a wide range of different elicitors and effectors. Some of these responses integrate into common downstream signalling pathways and regulate common sets of genes, while others are specific^89^. Therefore, RBPs participating in the response to flg22-sensing may just be the tip of the iceberg of the immune RBPome. On the other hand, pathogens develop mechanisms to manipulate the RBPome in order to create an infection favourable environment, as exemplified by GRP7, which is hijacked by the *P. syringae* effector HopU1^90^. Hence, RBPs can also be considered as susceptibility factors. Therefore, applying ptRIC to plants treated with a variety of elicitors, effectors, pathogens and hormones, will help to disentangle the networks of RBPs that work together during different immunity responses or that are targeted by pathogens.

Our results support the existence of ‘RBP-mediated plant immunity’, whereby RBPs orchestrate the reprogramming of RNA metabolism that occur during immune responses. Identification of the complement of RBPs that respond to flg22 represent a step forward towards understanding this reprogramming and provides an invaluable resource for the RNA, plant, immunology and microbiology communities. Moreover, this comparative ptRIC analysis can now be extended to virtually any physiological and pathological cue, which will likely expand our understanding on how gene expression adapts to an ever-changing environment.

This study has demonstrated that ptRIC is a suitable approach for the discovery of RBPs in plant tissues with important roles in immunity. Many of the flg22-responsive RBPs identified are also involved in responses to abiotic stresses, suggesting that they are promising multi-stress regulatory proteins. Therefore, our discoveries provide the foundations for the exploitation of knowledge of RBPs to develop crops with durable resistance to pathogens and other stresses.

## MATERIALS AND METHODS

### Plant material, treatment and plant RNA-interactome capture (ptRIC)

*Arabidopsis thaliana* plants were grown at neutral day conditions (12 h light, 12 h dark) at 20 °C and light intensity of approximately 100 μmol/m^2^/s. The mutant lines used in this study were obtained as T-DNA insertional lines from Nottingham Arabidopsis Stock Centre (NASC; http://arabidopsis.info). The mutant and overexpression lines used are detailed in **Table S5**. For the immune elicitation treatments, Arabidopsis leaves of mature plants (5-6 weeks old) were infiltrated with either flg22 (1 μM; Anaspec) or H_2_O (mock) using a 1 ml blunt end syringe and leaf tissue was harvested at 2 and 12 h after treatment. Four biological replicates per treatment and time point were performed. ptRIC was performed as previously described^12^, with NoCL controls processed in parallel with the CL samples. Whole cell proteome (input) samples for each of the treatments and time points were taken before performing ptRIC.

### SDS-PAGE, silver staining and western blot

Proteins were separated by SDS-PAGE (10-12% acrylamide) and transferred onto a PVDF membrane using the semidry TransBlot system (BioRad). The membrane was blocked with either 5% skimmed milk or 5% BSA (Bovine Serum Albumin) in TBST (Tris-Buffered Saline + 0.1 % tween) and incubated overnight at 4°C with one of the following antibodies: anti-METK1-4 (Agrisera, (Agrisera, AS163148A), anti-HSC70 AS08371), anti-RH3 (AS132714) or anti-MAPK (Cell Signalling Technology #4370). The membrane was washed 3 times with TBST, incubated with secondary antibody coupled to HRP (ThermoScientific, 31460) for 1 h at room temperature, washed again with TBST and visualised using LAS4000 (GE healthcare) and SuperSignal West Femto substrate (ThermoScientific). Protein extraction for the MAPK phosphorylation assay was performed as described by Flury and colleagues^91^.

### Sample preparation and MS analyses

MS analyses were performed as previously described^12^ with minor modifications. The isolated RBPs (eluates ptRIC) and whole cell proteome samples (inputs ptRIC) were prepared for MS using the standard FASP (filter aided sample preparation) method^92^ as previously described^12^. However, the total proteome samples were incubated with benzonase (Merk) for 45 minutes at 4°C prior to FASP to digest nucleic acids. The acidified tryptic digests were desalted on home-made 2 disc C18 StageTips as described^93^. After elution from the StageTips, samples were dried using a vacuum concentrator (Eppendorf) and the peptides were taken up in 10 µL 0.1% formic acid solution.

Experiments were performed on an Orbitrap Elite instrument (Thermo)^94^ that was coupled to an EASY-nLC 1000 liquid chromatography (LC) system (Thermo). The LC was operated in the one-column mode. The analytical column was a fused silica capillary (75 µm × 35 cm) with an integrated PicoFrit emitter (New Objective) packed in-house with Reprosil-Pur 120 C18-AQ 1.9 µm resin (Dr. Maisch). The analytical column was encased by a column oven (Sonation) and attached to a nanospray flex ion source (Thermo). The column oven temperature was adjusted to 45 °C during data acquisition. The LC was equipped with two mobile phases: solvent A (0.1% formic acid, FA, in water) and solvent B (0.1% FA in acetonitrile, ACN). All solvents were of UPLC grade (Sigma-Aldrich). Peptides were directly loaded onto the analytical column with a maximum flow rate that would not exceed the set pressure limit of 980 bar (usually around 0.6 – 1.0 µL/min). Peptides were subsequently separated on the analytical column by running a 140 min gradient of solvent A and solvent B (start with 7% B; gradient 7% to 35% B for 120 min; gradient 35% to 100% B for 10 min and 100% B for 10 min) at a flow rate of 300 nl/min. The mass spectrometer was operated using Xcalibur software (version 2.2 SP1.48). The mass spectrometer was set in the positive ion mode. Precursor ion scanning was performed in the Orbitrap analyzer (FTMS; Fourier Transform Mass Spectrometry) in the scan range of m/z 300-1800 and at a resolution of 60000 with the internal lock mass option turned on (lock mass was 445.120025 m/z, polysiloxane)^95^. Product ion spectra were recorded in a data dependent fashion in the ion trap (ITMS) in a variable scan range and at a rapid scan rate. The ionization potential (spray voltage) was set to 1.8 kV. Peptides were analyzed using a repeating cycle consisting of a full precursor ion scan (3.0 × 10^6^ ions or 50 ms) followed by 15 product ion scans (1.0 × 10^4^ ions or 50 ms) where peptides are isolated based on their intensity in the full survey scan (threshold of 500 counts) for tandem mass spectrum (MS2) generation that permits peptide sequencing and identification. Collision induced dissociation (CID) energy was set to 35% for the generation of MS2 spectra. During MS2 data acquisition dynamic ion exclusion was set to 120 seconds with a maximum list of excluded ions consisting of 500 members and a repeat count of one. Ion injection time prediction, preview mode for the FTMS, monoisotopic precursor selection and charge state screening were enabled. Only charge states higher than 1 were considered for fragmentation.

RAW spectra were submitted to an Andromeda^96^ search in MaxQuant (1.5.3.30) using the default settings^97^. Label-free quantification and match-between-runs was activated^98^. The MS/MS spectra data were searched against the Uniprot *A. thaliana* reference database (UP000006548_3702.fasta, downloaded 02/01/2018, one protein per gene). All searches included a contaminants database search (as implemented in MaxQuant, 245 entries). The contaminants database contains known MS contaminants and was included to estimate the level of contamination. Andromeda searches allowed oxidation of methionine residues (16 Da) and acetylation of the protein N-terminus (42 Da) as dynamic modifications and the static modification of cysteine (57 Da, alkylation with iodoacetamide). Enzyme specificity was set to “Trypsin/P” with two missed cleavages allowed. The instrument type in Andromeda searches was set to Orbitrap and the precursor mass tolerance was set to ±20 ppm (first search) and ±4.5 ppm (main search). The MS/MS match tolerance was set to ±0.5 Da. The peptide spectrum match FDR and the protein FDR were set to 0.01 (based on target-decoy approach). Minimum peptide length was 7 amino acids. For protein quantification unique and razor peptides were allowed. Modified peptides were allowed for quantification. The minimum score for modified peptides was 40. Label-free protein quantification was switched on, and unique and razor peptides were considered for quantification with a minimum ratio count of 2. Retention times were recalibrated based on the built-in nonlinear time-rescaling algorithm. MS/MS identifications were transferred between LC-MS/MS runs with the “match between runs” option in which the maximal match time window was set to 0.7 min and the alignment time window set to 20 min. The quantification is based on the “value at maximum” of the extracted ion current. At least two quantitation events were required for a quantifiable protein.

### Statistical analysis of the leaf RBPome

The MS data was analysed as previously described^12^ with minor modifications. After filtering for contaminants, protein groups that had at least one missing value in all biological conditions were filtered out to ensure we identified *bona fide* RBPs with consistent changes in association with RNA. After filtering, raw intensities were normalized using the variance stabilization normalization method implemented in the R/Bioconductor package ‘vsn’^99^. Then the pipeline of analysis continued as previously described^12^.

For the RBPs identified in ptRIC eluates, we identified batch effects that matched the dates when plant treatments and sample preparation were performed. These batch effects were accounted for by incorporating batch as block variable in linear modelling as described in the limma user guide^100^. However, for the ptRIC inputs (whole cell proteome) experiment the sample preparation was performed all together and the batch effect was minor. Thus, we decided not to incorporate ‘Batch’ into our linear model for the inputs. One of the replicates corresponding to the eluate of mock 12 hpt CL was lost during processing. Therefore, for that condition, the statistical analyses were performed using three biological samples. For both datasets (eluates and inputs), some of the identified peptides were not unique and could not be assigned to a single protein. For these, we performed the analysis using the protein groups. The protein groups are specified in the main text.

For the ptRIC eluate dataset, we defined proteins as RBPs based on statistical enrichment in CL over NoCL samples. For proteins with intensity values in both CL and NoCL samples, the log_2_ fold change between CL and NoCL samples (log_2_FC [CL/NoCL]) was calculated and moderate t-test and false discovery rate estimation (FDR) was used to calculate the statistical enrichment of CL/NoCL. We classified proteins as RBPs when FDR ≤ 0.01 and log_2_FC [CL/NoCL] ≥ 1.5 or FDR ≤ 0.1 and log_2_FC [CL/NoCL] ≥ 3.3. Proteins detected in none of the NoCL negative control samples and with intensity values in all the CL samples were also considered RBPs using a modification of the semiquantitative method as described previously^12^.

The identified RBPs were considered as ‘flg22-responsive’ when the differences between flg22 and mock treated samples were statistically significant. Flg22-responsive RBPs were classified into two different categories according to their statistical significance: FDR ≤ 0.1 (flg22-responsive RBPs) and 0.1 ≤ FDR ≤ 0.2 (candidate RBPs). Proteins that displayed significant differences but were not assigned RNA-binding activity at the condition with the highest intensity value were filtered out, i.e., flg22-stimulated RBPs had to be classified as RBPs at least in flg22-treated samples and flg22-inhibited RBPs had to be classified as RBPs at least in the mock-treated samples. For the ptRIC inputs dataset, proteins were considered as changing in abundance when differences between flg22 and mock treated samples identified with an FDR ≤ 0.1.

For the study of UV-crosslinkability, ptRIC eluates and input intensities were normalized using vsn method^99^, missing values were imputed by deterministic global minimal value using package ‘imputeLCMD’^101^, and processed intensities were proceeded to empirical Bayesian method moderated t-test using R package “limma”^100^. P values were adjusted for large sample size using Benjamini-Hochberg method.

For the clustering analysis, RBPs determined in ptRIC eluate experiments were divided into clusters based on their initial response to flg22, which is defined as fold change from mock to flg22 2 hpt, and progressive response to flg22, which is defined as fold change from flg22 2 hpt to flg22 12 hpt. Two stages of changes were plotted in scatter plot. Groups of dynamic behaviours were determined by both their positions in scatter plot, and moderated t-test result in ptRIC eluates. The cut-off line for t-test statistics was set at the same as flg22-responsive RBPs, cut-off lines for position in scatter plot were set at the minimal fold change in responsive RBPs.

### PTM enrichment

Annotation information for 15 different PTM types was downloaded from the Plant PTM Viewer database^22^. Enrichment of PTM occurrence was estimated based on hypergeometric distribution using R^102^. For each type of PTM, occurrence and p-value were calculated for its enrichment in flg22-responsive RBPs in each biological condition over the total RBPs detected. Relative differences in PTM annotations were log-transformed prior to calculate fold change.

### Data visualization

Graphs were generated using the ggplot2 package within R^103^. The Venn diagram was generated using the limma package within R^100^. To visualise the RBPomes in each of the conditions the missing values were imputed as described previously^12^ and all the proteins classified as RBPs (by both quantitative and semiquantitative) were included in the Volcano plot.

STRING analyses^29^ were performed to visualise the interactions between the flg22-responsive RBPs. The following parameters were used: meaning of network edges – confidence; active interaction sources: textmining, experiments and databases; minimum required interaction score: medium confidence.

The links to RNA biology of the identified RBPs were determined based on GO annotation as described by Beckmann and colleagues^104^.

### Prediction of the RNA-binding regions

The sequence propensity to bind RNA was calculated as described previously^87^.

### Genotyping of Arabidopsis mutants

To confirm the homozygous presence of the T-DNA insertion in both alleles, dual PCR was performed as detailed by O’Malley and colleagues^105^. Briefly, gDNA was extracted from the mutants and two independent PCRs were performed using different pairs of specific primers. The first PCR (T-DNA PCR) uses one primer located in the left border of the T-DNA (LB) and another primer in the genomic region at the 3’ end of the predicted insertion point (RP; **Table S6**). Thus, this PCR selectively amplifies a region that expands from the T-DNA to the specific genomic location within the gene. The second PCR (genomic PCR) uses a pair of primers designed in the genomic region around the T-DNA insertion point (LP and RP; **Table S6**).

### Bacterial growth assays

Leaves of mature Arabidopsis plants (5-6-week old) were syringe-infiltrated with *Pst*DC3000 or hrpA*-* at an inoculum density of 5×10^5^ cells/ml. At 0, 2 and 3 days after inoculation the bacterial density was analysed by homogenising leaf discs in water and plating serial dilutions onto LB (Luria-Bertani) plates with the appropriate antibiotics.

### Measurement of reactive oxygen species (ROS)

Measurements of ROS production were performed as described by Bach-Pages and Preston^106^ with minor modifications. Briefly, leaf discs of 4 mm diameter were punched from fully expanded leaves of mature plants using a cork borer and floated in water overnight at room temperature. Next day, leaf discs were transferred to a 96 well plate and presented with 200 μl of an elicitor solution containing HRP (20 μg/ml; ThermoFisher), LO-12 (10 μM; WAKO) and flg22 (500 nM; Anaspec). Luminescence was measured every minute using a M1000Pro microplate reader (TECAN Group Ltd.) for 1 h and integration time 200 ms.

## ACKNOWLEDGEMENTS

We want to thank all the members of Preston, Castello and van der Hoorn lab for their fruitful discussions. We thank Urszula Pyzio, Sarah Rodgers and Caroline O’Brien for excellent technical support. We thank Hong Qiao for the EIN2OE seeds. Marcel Bach-Pages is supported by Biotechnology and Biological Sciences Research Council (BBSRC, grant BB/M011224/1), Lorna Casselton Memorial Scholarship at St. Cross College (University of Oxford) and a John Fell Fund (University of Oxford) award to Gail Preston and Alfredo Castello. Renier van der Hoorn is supported by ERC grant 616449 ‘GreenProteases’. Alfredo Castello is supported by MRC Career Development Award MR/L019434/1 and MRC grant MR/R021562/1.

## AUTHOR CONTRIBUTIONS

MBP, RvdH, AC and GMP designed the experiments. MBP performed all the experiments except for the proteomic experiment, which was performed by FK and MK. HC, NS, RS and SM assisted with the bioinformatics analyses. MBP, GMP and AC analysed the data and wrote the manuscript.

## SUPPLEMENTARY DATA

### Supplementary Figures 1 - 6

**Fig. S1**. Arabidopsis high confidence leaf RBPomes

**Fig. S2**. Response of the whole cell proteome (input) to flg22 perception

**Fig. S3**. Changes in association with RNA of the RBPome are not primarily driven by changes in protein abundance

**Fig. S4**. RBP networks altered during flg22-induced plant immune response

**Fig. S5**. Mutant lines used for phenotypic mutant screen

**Fig. S6**. Mode of action of EIN2

### Supplementary Tables

**Table S1**. Arabidopsis leaf RBPome

**Table S2**. Flg22-responsive RBPome

**Table S3**. Total proteome (WCL) responses to flg22

**Table S4**. Cross-reference of input (total protein) and eluate (RNA-bound fraction) data

**Table S5**. Arabidopsis lines screened

**Table S6**. List of primers used in this study

## REFERENCES

1. Zipfel, C. Plant pattern-recognition receptors. Trends Immunol. 35, 345–351 (2014).

2. Chiang, Y.-H. & Coaker, G. Effector triggered immunity: NLR immune perception and downstream defense responses. Arab. B. 13, e0183 (2015).

3. Andersen, E. J., Ali, S., Byamukama, E., Yen, Y. & Nepal, M. P. Disease Resistance Mechanisms in Plants. Genes (Basel). 9, 339 (2018).

4. Yu, X. et al. Orchestration of processing body dynamics and mRNA decay in Arabidopsis immunity. Cell Rep. 28, 2194-2205.e6 (2019).

5. Glisovic, T., Bachorik, J. L., Yong, J. & Dreyfuss, G. RNA-binding proteins and post-transcriptional gene regulation. FEBS Lett. 582, 1977–1986 (2008).

6. Staiger, D., Korneli, C., Lummer, M. & Navarro, L. Emerging role for RNA-based regulation in plant immunity. New Phytol. 197, 394–404 (2013).

7. Castello, A. et al. Insights into RNA biology from an atlas of mammalian mRNA-binding proteins. Cell 149, 1393–1406 (2012).

8. Baltz, A. G. et al. The mRNA-bound proteome and its global occupancy profile on protein-coding transcripts. Mol. Cell 46, 674–690 (2012).

9. Hentze, M. W., Castello, A., Schwarzl, T. & Preiss, T. A brave new world of RNA-binding proteins. Nat. Rev. Mol. Cell Biol. 19, 327–341 (2018).

10. Garcia-Moreno, M. et al. System-wide profiling of RNA-binding proteins uncovers key regulators of virus infection. Mol. Cell 74, 1–16 (2019).

11. Sysoev, V. O. et al. Global changes of the RNA-bound proteome during the maternal-to-zygotic transition in Drosophila. Nat. Commun. 7, 12128 (2016).

12. Bach-Pages, M. et al. Discovering the RNA-binding proteome of plant leaves with an improved RNA interactome capture method. Biomolecules 10, 661 (2020).

13. Reichel, M. et al. In planta determination of the mRNA-binding proteome of Arabidopsis etiolated seedlings. Plant Cell 28, 2435–2452 (2016).

14. Marondedze, C., Thomas, L., Serrano, N. L., Lilley, K. S. & Gehring, C. The RNA-binding protein repertoire of *Arabidopsis thaliana*. Sci. Rep. 6, 29766 (2016).

15. Zhang, Z. et al. UV crosslinked mRNA-binding proteins captured from leaf mesophyll protoplasts. Plant Methods 12, 42 (2016).

16. Marondedze, C., Thomas, L., Gehring, C. & Lilley, K. S. Changes in the Arabidopsis RNA-binding proteome reveal novel stress response mechanisms. BMC Plant Biol. 19, 1–11 (2019).

17. Bach-Pages, M., Castello, A. & Preston, G. M. Plant RNA interactome capture: revealing the plant RBPome. Trends Plant Sci. 22, 449–451 (2017).

18. Suarez Rodriguez, M. C., Petersen, M. & Mundy, J. Mitogen-activated protein kinase signaling in plants. Annu. Rev. Plant Biol. 61, 621–649 (2010).

19. Li, Y. et al. Identification of microRNAs involved in pathogen-associated molecular pattern-triggered plant innate immunity. Plant Physiol. 152, 2222–2231 (2010).

20. Castello, A. et al. Comprehensive identification of RNA-binding domains in human cells. Mol. Cell 63, 696–710 (2016).

21. Arif, A. et al. The GAIT translational control. Wiley Interdiscip Rev RNA 9, 1441 (2018).

22. Willems, P. et al. The Plant PTM Viewer, a central resource for exploring plant protein modifications. Plant J. 99, 752–762 (2019).

23. Cui, B. et al. S-nitrosylation of the zinc finger protein SRG1 regulates plant immunity. Nat. Commun. 9, 1–12 (2018).

24. Marino, D., Peeters, N. & Rivas, S. Ubiquitination during plant immune signaling. Plant Physiol. 160, 15–27 (2012).

25. Garcia-Moreno, M. et al. System-wide Profiling of RNA-Binding Proteins Uncovers Key Regulators of Virus Infection. Mol. Cell 74, 196-211.e11 (2019).

26. Denoux, C. et al. Activation of defense response pathways by OGs and Flg22 elicitors in Arabidopsis seedlings. Mol. Plant 1, 423–445 (2008).

27. Kilchert, C. et al. System-wide analyses of the fission yeast poly(A)+ RNA interactome reveal insights into organization and function of RNA–protein complexes. Genome Res. 1–15 (2020). doi: 10.1101/gr.257006.119

28. Castello, A., Hentze, M. W. & Preiss, T. Metabolic enzymes enjoying new partnerships as RNA-binding proteins. Trends Endocrinol. Metab. 26, 746–757 (2015).

29. Szklarczyk, D. et al. The STRING database in 2017: Quality-controlled protein-protein association networks, made broadly accessible. Nucleic Acids Res. 45, D362–D368 (2017).

30. Entwisle, S. W. et al. Proteome and Phosphoproteome Analysis of Brown Adipocytes Reveals That RICTOR Loss Dampens Global Insulin/AKT Signaling. Mol. Cell. Proteomics 19, 1104–1119 (2020).

31. Sun, T. et al. An RNA recognition motif-containing protein is required for plastid RNA editing in Arabidopsis and maize. Proc. Natl. Acad. Sci. U. S. A. 110, 1169–1178 (2013).

32. Næsted, H. et al. Arabidopsis VARIEGATED 3 encodes a chloroplast-targeted, zinc-finger protein required for chloroplast and palisade cell development. J. Cell Sci. 117, 4807–4818 (2004).

33. García-Andrade, J., Ramírez, V., López, A. & Vera, P. Mediated Plastid RNA Editing in Plant Immunity. PLoS Pathog. 9, 1–13 (2013).

34. Xu, G. et al. Global translational reprogramming is a fundamental layer of immune regulation in plants. Nature 545, 487–490 (2017).

35. Yoo, H. et al. Translational Regulation of Metabolic Dynamics during Effector-Triggered Immunity. Mol. Plant 88–98 (2019). doi: 10.1016/j.molp.2019.09.009

36. Meteignier, L. V. et al. Translatome analysis of an NB-LRR immune response identifies important contributors to plant immunity in Arabidopsis. J. Exp. Bot. 68, 2333–2344 (2017).

37. Machado, J. P. B., Calil, I. P., Santos, A. A. & Fontes, E. P. B. Translational control in plant antiviral immunity. Genet. Mol. Biol. 40, 292–304 (2017).

38. Mustroph, A. et al. Profiling translatomes of discrete cell populations resolves altered cellular priorities during hypoxia in Arabidopsis. Proc. Natl. Acad. Sci. U. S. A. 106, 18843–18848 (2009).

39. Yángüez, E., Castro-Sanz, A. B., Fernández-Bautista, N., Oliveros, J. C. & Castellano, M. M. Analysis of genome-wide changes in the translatome of Arabidopsis seedlings subjected to heat stress. PLoS One 8, e71425 (2013).

40. Lei, L. et al. Ribosome profiling reveals dynamic translational landscape in maize seedlings under drought stress. Plant J. 84, 1206–1208 (2015).

41. Ross, S., Giglione, C., Pierre, M., Espagne, C. & Meinnel, T. Functional and developmental impact of cytosolic protein N-terminal methionine excision in Arabidopsis. Plant Physiol. 137, 623–637 (2005).

42. Ree, R., Varland, S. & Arnesen, T. Spotlight on protein N-terminal acetylation. Exp. Mol. Med. 50, 90 (2018).

43. Xu, F. et al. Two N-terminal acetyltransferases antagonistically regulate the stability of a nod-like receptor in arabidopsis. Plant Cell 27, 1547–1562 (2015).

44. Boisson, B., Giglione, C. & Meinnel, T. Unexpected protein families including cell defense components feature in the N-myristoylome of a higher eukaryote. J. Biol. Chem. 278, 43418–43429 (2003).

45. Ohta, H., Takamune, N., Kishimoto, N., Shoji, S. & Misumi, S. N-myristoyltransferase 1 enhances immunodeficiency human virus replication through regulation of viral RNA expression level. Biochem. Biophys. Res. Commun. 463, 988–993 (2015).

46. Chantarachot, T. & Bailey-Serres, J. Polysomes, stress granules, and processing bodies: A dynamic triumvirate controlling cytoplasmic mRNA fate and function. Plant Physiol. 176, 254–269 (2018).

47. Weber, C., Nover, L. & Fauth, M. Plant stress granules and mRNA processing bodies are distinct from heat stress granules. Plant J. 56, 517–530 (2008).

48. Maldonado-Bonilla, L. D. et al. The arabidopsis tandem zinc finger 9 protein binds RNA and mediates pathogen-associated molecular pattern-triggered immune responses. Plant Cell Physiol. 55, 412–425 (2014).

49. Tharun, S. Lsm1-7-Pat1 complex: a link between 3’ and 5’-ends in mRNA decay? RNA Biol. 6, 228–232 (2009).

50. Roux, M. E. et al. The mRNA decay factor PAT1 functions in a pathway including MAP kinase 4 and immune receptor SUMM2. EMBO J. 34, 593–608 (2015).

51. Chantarachot, T. et al. DHH1/DDX6-like RNA helicases maintain ephemeral half-lives of stress-response mRNAs. Nature Plants 6, 675–685 (2020).

52. Horos, R. et al. The small non-coding vault RNA1-1 acts as a riboregulator of autophagy. Cell 176, 1054-1067.e12 (2019).

53. Choudhury, N. R. et al. RNA-binding activity of TRIM25 is mediated by its PRY/SPRY domain and is required for ubiquitination. BMC Biol. 15, 1–20 (2017).

54. Thapar, R. Roles of prolyl isomerases in RNA-mediated gene expression. Biomolecules 5, 974–999 (2015).

55. Vijayapalani, P., Hewezi, T., Pontvianne, F. & Baum, T. J. An effector from the cyst nematode heterodera schachtii derepresses host rRNA genes by altering histone acetylation. Plant Cell 30, 2795–2812 (2018).

56. Park, C. J. & Seo, Y. S. Heat shock proteins: a review of the molecular chaperones for plant immunity. Plant Pathol. J. 31, 323–333 (2015).

57. Lopez, V. A. et al. A Bacterial Effector Mimics a Host HSP90 Client to Undermine Immunity. Cell 179, 205-218.e21 (2019).

58. Järvelin, A. I., Noerenberg, M., Davis, I. & Castello, A. The new (dis) order in RNA regulation. Cell Commun. Signal. 14, (2016).

59. Datta, R. et al. Glutathione regulates 1-aminocyclopropane-1-carboxylate synthase transcription via WRKY33 and 1-aminocyclopropane-1-carboxylate oxidase by modulating messenger RNA stability to induce ethylene synthesis during stress. Plant Physiol. 169, 2963–2981 (2015).

60. Xu, Y., Wang, B. C. & Zhu, Y. X. Identification of proteins expressed at extremely low level in Arabidopsis leaves. Biochem. Biophys. Res. Commun. 358, 808–812 (2007).

61. Chen, C. Y. A. & Shyu, A. Bin. Emerging mechanisms of mRNP remodeling regulation. Wiley Interdiscip. Rev. RNA 5, 713–722 (2014).

62. Li, D., Liu, H., Zhang, H., Wang, X. & Song, F. OsBIRH1, a DEAD-box RNA helicase with functions in modulating defence responses against pathogen infection and oxidative stress. J. Exp. Bot. 59, 2133–2146 (2008).

63. Chaudhary, S. et al. Alternative splicing and protein diversity: plants versus animals. Front. Plant Sci. 10, 1–14 (2019).

64. Feng, J. et al. SKIP confers osmotic tolerance during salt stress by controlling alternative gene splicing in Arabidopsis. Mol. Plant 8, 1038–1052 (2015).

65. Cohen, I., Sapir, Y. & Shapira, M. A conserved mechanism controls translation of Rubisco large subunit in different photosynthetic organisms. 141, 1089–1097 (2006).

66. Choquet, Y., Zito, F., Wostrikoff, K. & Wollman, F. A. Cytochrome f translation in *Chlamydomonas* chloroplast is autoregulated by its carboxyl-terminal domain. Plant Cell 15, 1443–1454 (2003).

67. Müller-Mcnicoll, M., Rossbach, O., Hui, J. & Medenbach, J. Auto-regulatory feedback by RNA-binding proteins. J. Mol. Cell Biol. 11, 930–939 (2019).

68. Wei, W. et al. The gene coding for the Hrp pilus structural protein is required for type III secretion of Hrp and Avr proteins in Pseudomonas syringae pv. tomato. Proc. Natl. Acad. Sci. U. S. A. 97, 2247–2252 (2000).

69. Mnatsakanyan, R. et al. Proteome-wide detection of S-nitrosylation targets and motifs using bioorthogonal cleavable-linker-based enrichment and switch technique. Nat. Commun. 10, 1–12 (2019).

70. Chicois, C. et al. The UPF1 interactome reveals interaction networks between RNA degradation and translation repression factors in Arabidopsis. Plant J. 96, 119–132 (2018).

71. Koncz, C., DeJong, F., Villacorta, N., Szakonyi, D. & Koncz, Z. The spliceosome-activating complex: molecular mechanisms underlying the function of a pleiotropic regulator. Front. Plant Sci. 3, 1–12 (2012).

72. Liu, Y. & Imai, R. Function of plant DExD/H-Box RNA helicases associated with ribosomal RNA biogenesis. Front. Plant Sci. 9, 1–7 (2018).

73. Sauter, M., Moffatt, B., Saechao, M. C., Hell, R. & Wirtz, M. Methionine salvage and S-adenosylmethionine: Essential links between sulfur, ethylene and polyamine biosynthesis. Biochem. J. 451, 145–154 (2013).

74. Meng, J. et al. METHIONINE ADENOSYLTRANSFERASE4 mediates DNA and histone methylation. Plant Physiol. 177, 652–670 (2018).

75. Zhou, W., Karcher, D. & Bock, R. Identification of enzymes for adenosine-to-inosine editing and discovery of cytidine-to-uridine editing in nucleus-encoded transfer RNAs of arabidopsis. Plant Physiol. 166, 1985–1997 (2014).

76. Ju, C. et al. Conservation of ethylene as a plant hormone over 450 million years of evolution. Nat. Plants 1, 1–7 (2015).

77. Rin, S. et al. EIN2-mediated signaling is involved in pre-invasion defense in Nicotiana benthamiana against potato late blight pathogen, Phytophthora infestans. Plant Signal. Behav. 12, e1300733 (2017).

78. Alonso, J. M., Hirayama, T., Roman, G., Nourizadeh, S. & Ecker, J. R. EIN2, a bifunctional transducer of ethylene and stress responses in Arabidopsis. Science (80-.). 284, 2148–2152 (1999).

79. Tintor, N. et al. Layered pattern receptor signaling via ethylene and endogenous elicitor peptides during Arabidopsis immunity to bacterial infection. Proc Natl Acad Sci U S A 110, 6211–6216 (2013).

80. Mersmann, S., Bourdais, G., Rietz, S. & Robatzek, S. Ethylene signaling regulates accumulation of the FLS2 receptor and is required for the oxidative burst contributing to plant immunity. Plant Physiol. 154, 391–400 (2010).

81. Boutrot, F. et al. Direct transcriptional control of the Arabidopsis immune receptor FLS2 by the ethylene-dependent transcription factors EIN3 and EIL1. Proc. Natl. Acad. Sci. U. S. A. 107, 14502–14507 (2010).

82. Ju, C. et al. CTR1 phosphorylates the central regulator EIN2 to control ethylene hormone signaling from the ER membrane to the nucleus in Arabidopsis. PNAS 109, 19486–19491 (2012).

83. Qiao, H. et al. Processing and subcellular trafficking of ER-tethered EIN2 control response to ethylene gas. Science 338, 390–393 (2012).

84. Wen, X. et al. Activation of ethylene signaling is mediated by nuclear translocation of the cleaved EIN2 carboxyl terminus. Cell Res. 22, 1613–1616 (2012).

85. Li, W. et al. EIN2-directed translational regulation of ethylene signaling in Arabidopsis. Cell 163, (2015).

86. Merchante, C. et al. Gene-specific translation regulation mediated by the hormone-signaling molecule EIN2. Cell 163, 684–697 (2015).

87. Hobor, F. et al. A cryptic RNA-binding domain mediates Syncrip recognition and exosomal partitioning of miRNA targets. Nat. Commun. 9, (2018).

88. Köster, T. & Meyer, K. Plant ribonomics: proteins in search of RNA partners. Trends Plant Sci. 23, 352–365 (2018).

89. Buscaill, P. & Rivas, S. Transcriptional control of plant defence responses. Curr. Opin. Plant Biol. 20, 35–46 (2014).

90. Fu, Z. Q. et al. A type III effector ADP-ribosylates RNA-binding proteins and quells plant immunity. Nature 447, 284–288 (2007).

91. Flury, P., Klauser, D., Boller, T. & Bartels, S. MAPK phosphorylation assay with leaf disks of Arabidopsis. Bio-protocol 3, 3–6 (2013).

92. Wiśniewski, J. R., Zougman, A., Nagaraj, N. & Mann, M. Universal sample preparation method for proteome analysis. Nat. Methods 6, 359–62 (2009).

93. Rappsilber, J., Mann, M. & Ishihama, Y. Protocol for micro-purification, enrichment, pre-fractionation and storage of peptides for proteomics using StageTips. Nat. Protoc. 2, 1896–1906 (2007).

94. Michalski, A. et al. Ultra high resolution linear ion trap orbitrap mass spectrometer (orbitrap elite) facilitates top down LC MS/MS and versatile peptide fragmentation modes. Mol. Cell. Proteomics 11, 1–11 (2012).

95. Olsen, J. V. et al. Parts per million mass accuracy on an orbitrap mass spectrometer via lock mass injection into a C-trap. Mol. Cell. Proteomics 4, 2010–2021 (2005).

96. Cox, J. et al. Andromeda: a peptide search engine integrated into the MaxQuant environment. J. Proteome Res. 10, 1794–1805 (2011).

97. Cox, J. & Mann, M. MaxQuant enables high peptide identification rates, individualized p.p.b.-range mass accuracies and proteome-wide protein quantification. Nat. Biotechnol. 26, 1367–1372 (2008).

98. Cox, J. et al. Accurate proteome-wide label-free quantification by delayed normalization and maximal peptide ratio extraction, termed MaxLFQ. Mol. Cell. Proteomics 13, 2513–2526 (2014).

99. Huber, W., Poustka, A. & Vingron, M. Variance stabilization applied to microarray data calibration and to the quantification of differential expression. Bioinformatics 18, S96–S104 (2002).

100. Smyth, G. K. Linear models and empirical bayes methods for assessing differential expression in microarray experiments. Stat. Appl. Genet. Mol. Biol. 3, 1–25 (2004).

101. Package, T. & Lazar, A. C. A collection of methods for left-censored missing data imputation. 1–30 (2015).

102. R Core Team. R: A language and environment for statistical computing. R Foundation for Statistical Computing. (2017).

103. Wickham, H. ggplot2: Elegant Graphics for Data Analysis. ggplot2 - Elegant Graphics for Data Analysis (Springer-Verlag, 2016). doi: 10.18637/jss.v077.b02

104. Beckmann, B. M. et al. The RNA-binding proteomes from yeast to man harbour conserved enigmRBPs. Nat. Commun. 6, 10127 (2015).

105. O’Malley, R. C., Barragan, C. C. & Ecker, J. R. A user’s guide to the Arabidopsis T-DNA insertional mutant collections. Methods Mol. Biol. 1284, 323–342 (2015).

106. Bach-Pages, M. & Preston, G. M. Methods to quantify biotic-induced stress in plants. in Host-Pathogen Interactions 1734, 241–255 (2018).

