## Supplementary figures 1-6 for "Proteome-wide Profiling of RNA-Binding Protein Responses to flg22 Reveals Novel Components of Plant Immunity"

Figure S1
Figure S2
Figure S3
Figure S4
Figure S5
Figure S6

#### Figure S1

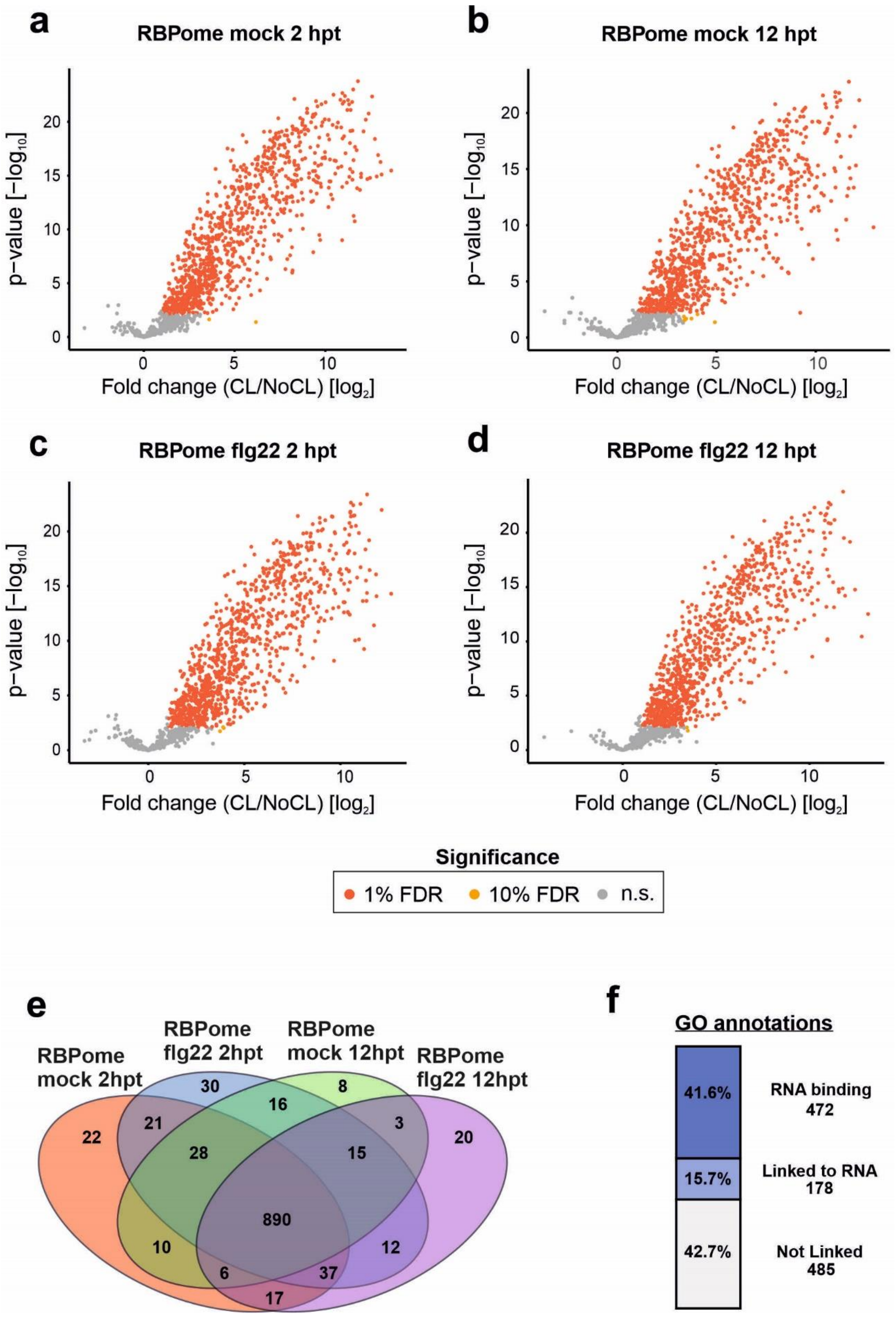

a
RBPome mock 2 hpt
p−value [−log10]
Fold change (CL/NoCL) [log2]
b
RBPome mock 12 hpt
c
RBPome flg22 2 hpt
d
RBPome flg22 12 hpt
Significance
1% FDR
10% FDR
n.s.
e
RBPome mock 2hpt
RBPome flg22 2hpt
RBPome mock 12hpt
RBPome flg22 12hpt
22
30
8
20
21
16
3
28
15
890
10
12
6
37
17
f
GO annotations
41.6%
RNA binding 472
15.7%
Linked to RNA 178
42.7%
Not Linked 485

**Fig.S1. Arabidopsis high confidence leaf RBPomes. a-d** Volcano plots depicting the $log_2$ fold change and the significance (p-value) of each protein (dots) between UV-crosslinking (CL) and non-crosslinked (NoCL) treatment using data from four biological replicates. Proteins are coloured in red when false discovery rate (FDR) $\leq 0.01$ and $log_2FC$ [CL/NoCL] $\geq 1.5$ and proteins are coloured in yellow when FDR $\leq 0.1$ and $log_2FC$ [CL/NoCL] $\geq 3.3$. Non-significant proteins are coloured in grey. Red and orange proteins represent the high confidence leaf RBPome. **e** Venn diagram showing the overlap between the four high confidence leaf RBPome datasets. **f** Proportion of high confidence leaf RBPs linked to RNA biology or not based on GO annotations.

### Figure S2

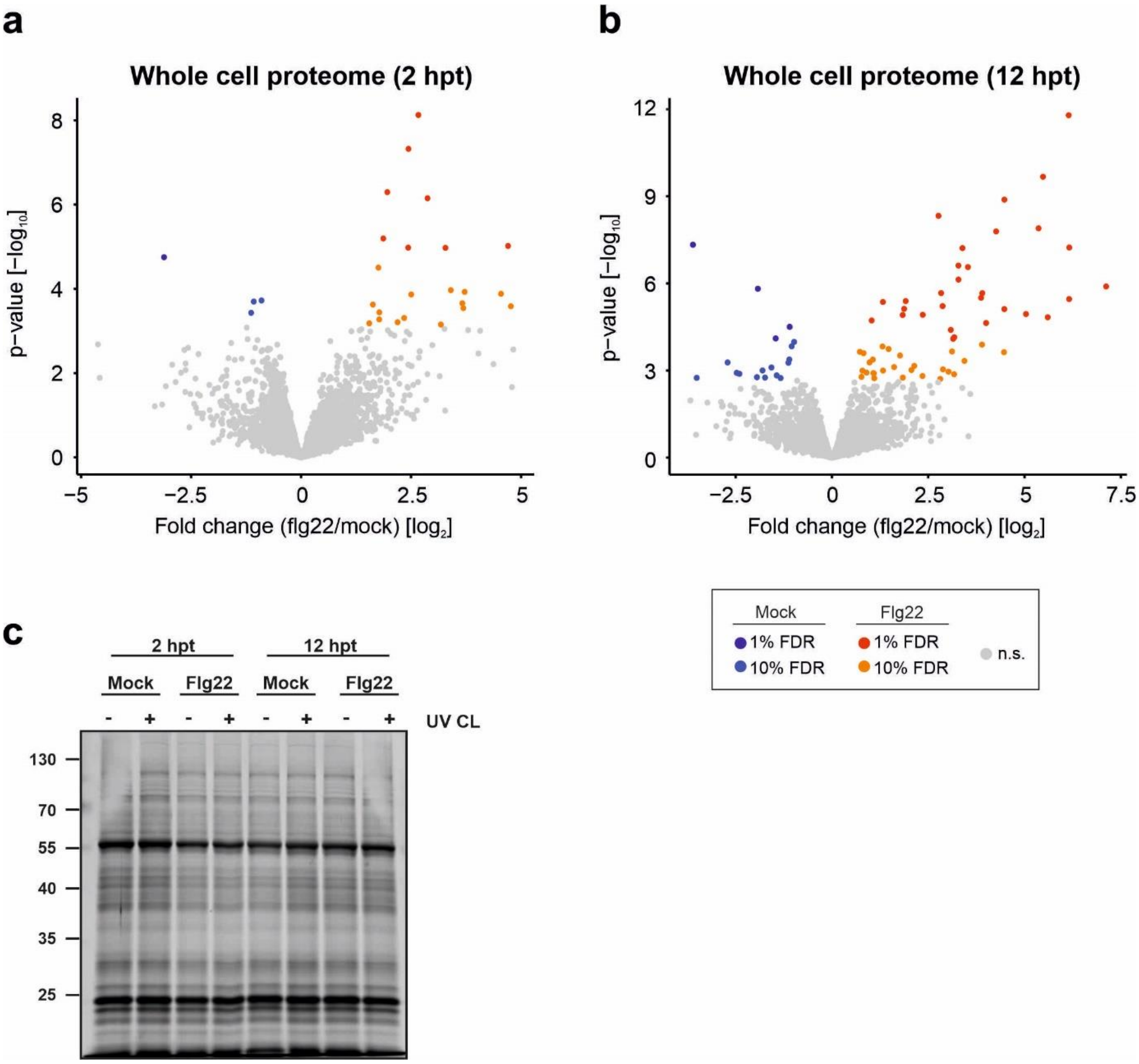


**Fig.S2. Response of the whole cell proteome (input) to flg22 perception. a-b** Volcano plots depicting the $\log_2$ fold change and the significance (p-value) of each protein (dots) between flg22 and mock treatment using data from four biological replicates. Proteins are coloured in red when false discovery rate (FDR) ≤ 0.01 and $\log_2$FC [flg22/mock] > 0, in orange when FDR ≤ 0.1 and $\log_2$FC [flg22/mock] > 0, in dark blue when FDR ≤ 0.01 and $\log_2$FC [flg22/mock] < 0 and in light blue when FDR ≤ 0.1 and $\log_2$FC [flg22/mock] < 0. Red and orange proteins represent proteins with higher accumulation upon flg22 perception, whereas dark blue and light blue represent proteins with lower accumulation upon flg22 perception. Non-significant proteins are coloured in grey. **c** Silver staining analyses of the whole cell proteomes (inputs). The whole cell proteomes of Arabidopsis leaves treated with either mock ($H_2O$) or flg22 were separated in a 12% acrylamide gel and analysed by silver staining

### Figure S3

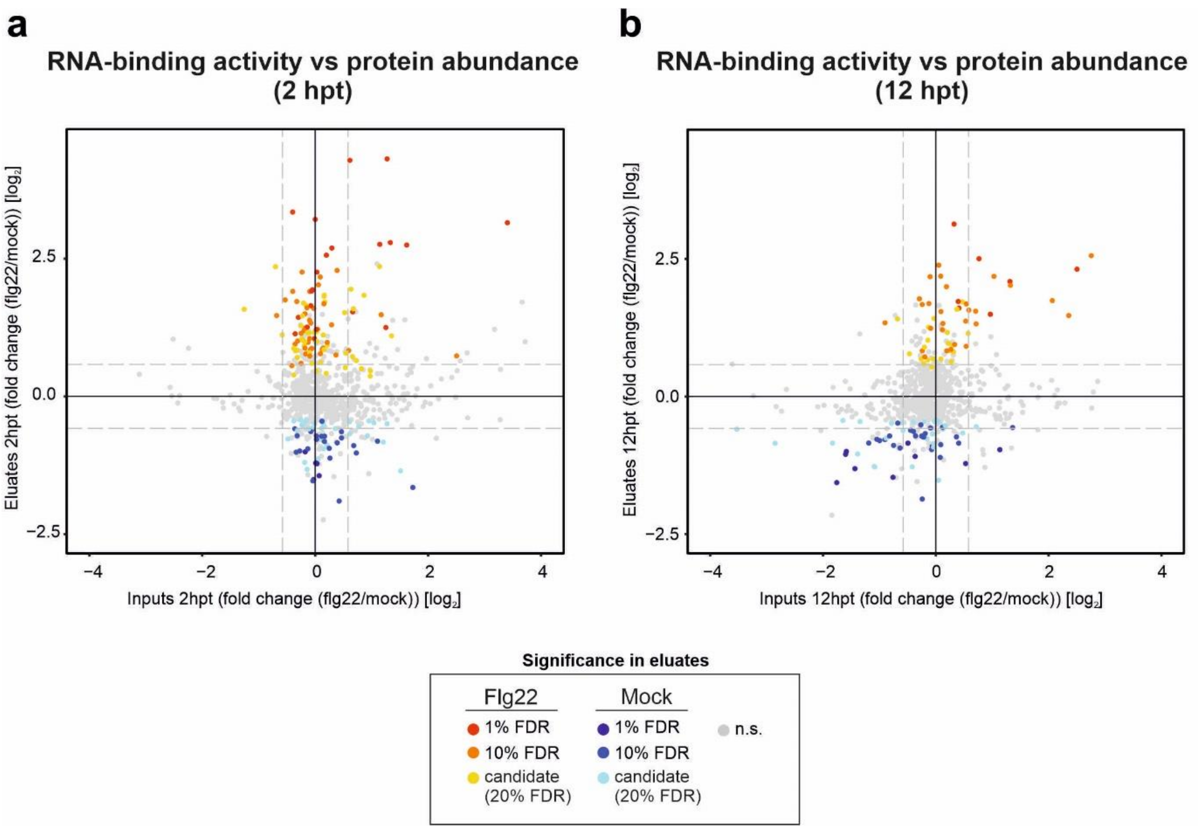


**Fig. S3. Changes in association with RNA of the RBPome are not primarily driven by changes in protein abundance.** Scatter plots depicting the $\log_2$ fold change in ptRIC inputs (x-axis) and $\log_2$ fold change in ptRIC eluates (y-axis) of each protein (dots) between flg22 and mock treatment using data from four biological replicates. Proteins are coloured according to significance (false discovery rate, FDR) in ptRIC eluates as following: proteins are coloured in red when FDR ≤ 0.01 and $\log_2$FC[flg22/mock] > 0, in orange when FDR ≤ 0.1 and $\log_2$FC [flg22/mock] > 0, in yellow when FDR ≤ 0.2 and $\log_2$FC [flg22/mock] > 0, in dark blue when FDR ≤ 0.01 and $\log_2$FC [flg22/mock] < 0, in blue when FDR ≤ 0.1 and $\log_2$FC [flg22/mock] < 0 and in light blue when FDR ≤ 0.2 and $\log_2$FC [flg22/mock] < 0. Red, orange and yellow proteins represent the leaf RBPs stimulated upon flg22 perception (ptRIC), whereas dark blue, blue and light blue represent leaf RBPs inhibited upon flg22 perception (ptRIC). Non-significant proteins in ptRIC are coloured in grey.

### Figure S4

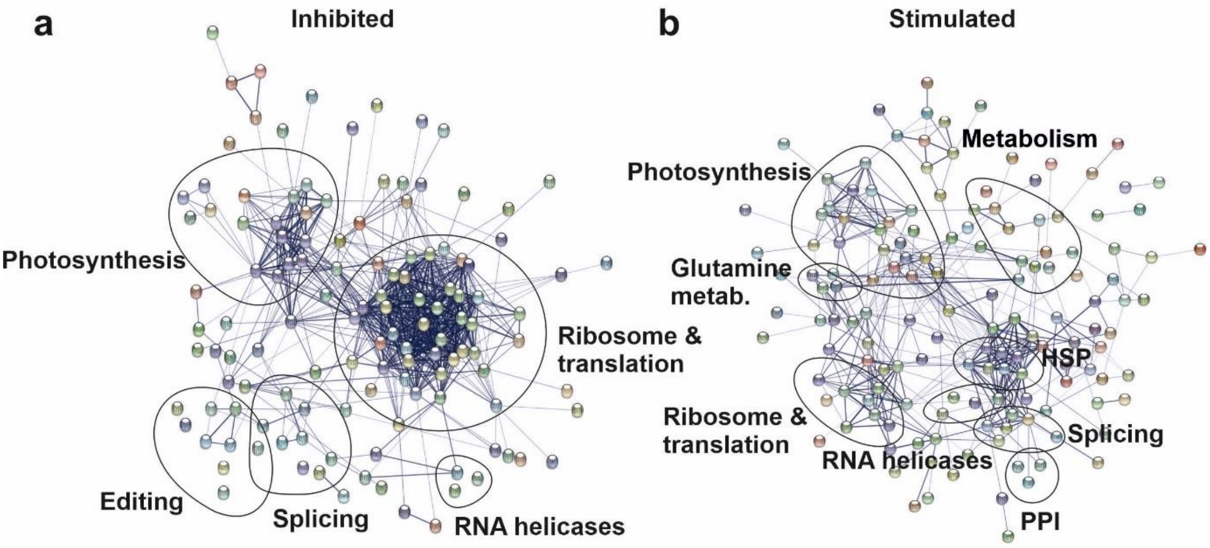


**Fig. S4. RBP networks altered during flg22-induced plant immune response.** STRING network depicting the protein-protein interactions between the flg22-inhibited and flg22-stimulated RBPs. Each dot represents a protein and the thickness of lines represents the confidence of the interactions.

### Figure S5

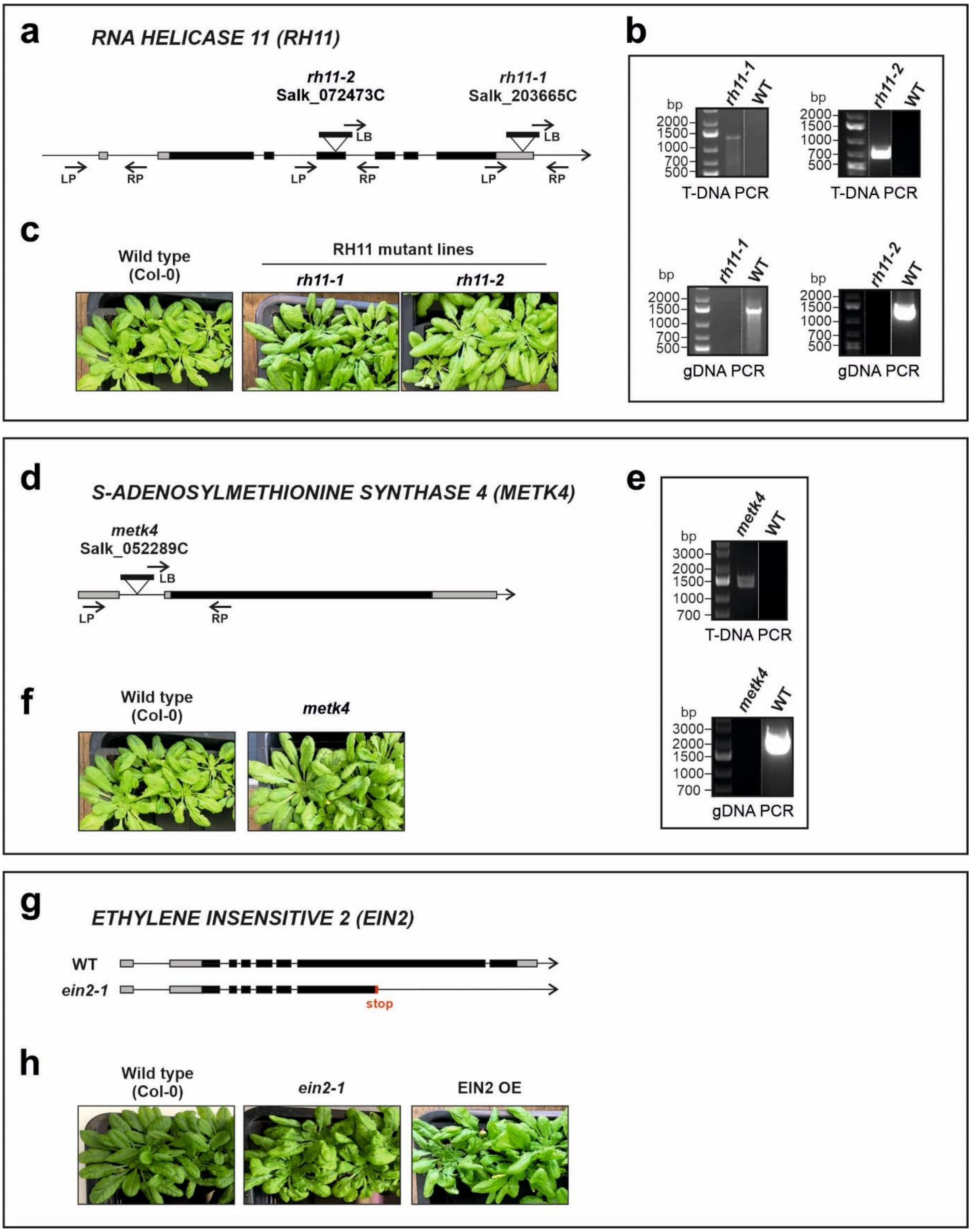


**Fig. S5. Mutant lines used for phenotypic mutant screen. a, d, g** Schematic representation of the insertion of T-DNA for each of the mutant lines for RH11 and METK4 and the *ein2-1* mutant. **b, e** Dual genotyping PCRs of the mutant lines: T-DNA PCR was performed using the

T-DNA LB (Salk) and gene-specific RP, whereas the gDNA was performed with gene-specific LP and RP. **c, f, h** Representative pictures of 6-week old plants of each mutant lines, overexpressing lines and Col-0 (WT).

### Figure S6

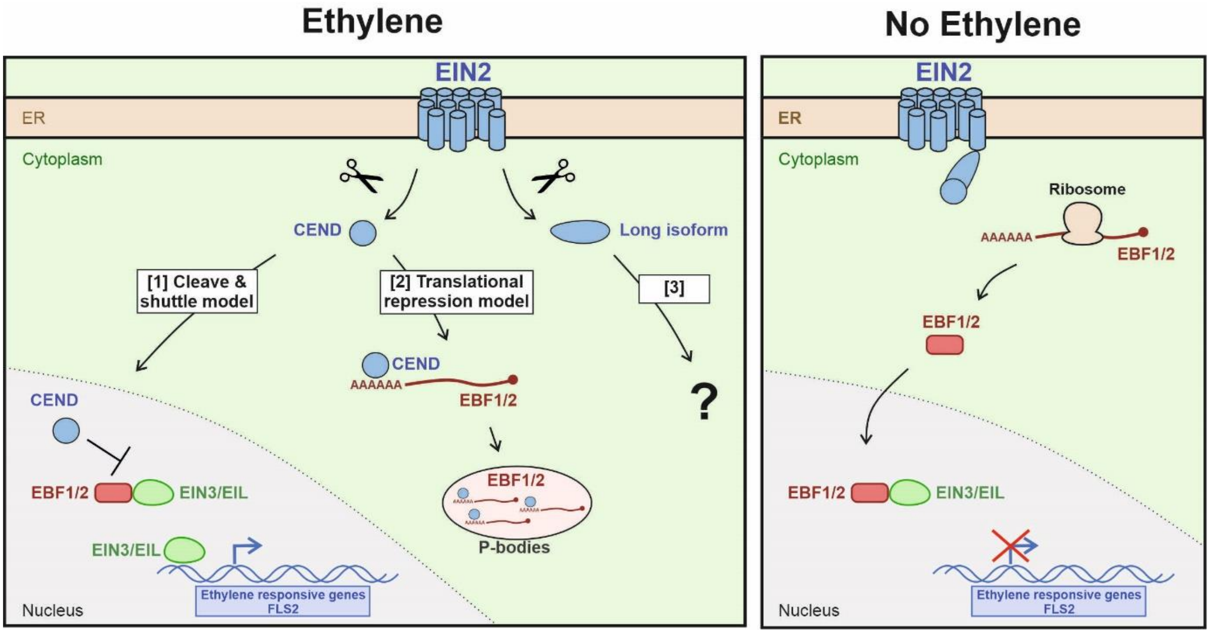


**Fig. S6. Mode of action of EIN2.** Upon ethylene perception, the CEND domain is cleaved and can play two different roles: [1] the CEND can shuttle to the nucleus to stabilize two transcription factors (EIN3/EIL1) that positively regulate ethylene responses; [2] the CEND can bind the 3'UTR of the negative regulators EBF1/2 in the cytoplasm and together with the NMD machinery re-localise them to the PBs to promote translational repression. EBF1/2 promotes degradation of EIN3/EIL1 transcription factors. Hence, translational repression of EBF1/2, results in the expression of ethylene responsive genes. [3] A longer isoform was identified in this study, however, its role and mechanism remain unknown.
